# IRIS-EDA: An integrated RNA-Seq interpretation system for gene expression data analysis

**DOI:** 10.1101/283341

**Authors:** Brandon Monier, Adam McDermaid, Jing Zhao, Anne Fennell, Qin Ma

## Abstract

**Motivation:** Next-Generation Sequencing has made available substantial amounts of large-scale genomic and transcriptomic data. Studies with RNA-Sequencing (RNA-Seq) data typically involve generation of gene expression profiles that can be further analyzed using correlation analysis, co-expression analysis, clustering, differential gene expression (DGE), among many other studies. While these analyses can provide invaluable information related to gene expression, interpretation of the results can prove challenging.

**Results:** Here we present a tool called IRIS-EDA, which is a web-server-based expression data analysis tool developed using Shiny. It provides a straightforward and user-friendly platform for performing numerous computational analyses (e.g., principal component analysis, clustering, biclustering, and DGE) on user-provided gene expression data generated from both bulk RNA-Seq and Single-cell RNA-Seq data. Specifically, three most commonly used R-based tools (edgeR, DESeq2, and limma) are implemented in the DGE analysis with seven unique experimental design functionalities, including a user-specified design matrix option. Six discovery-driven methods and tools (correlation analysis, heatmap, clustering, biclustering, PCA, and MDS) are provided for gene expression exploration which is useful for designing experimental hypotheses and determining key factors for comprehensive DGE analysis. Furthermore, this platform integrates seven visualization tools in a highly interactive manner, for improved interpretation of the analyses. It is noteworthy that, for the first time, IRIS-EDA provides a framework to expedite submission of data and results to NCBI’s Gene Expression Omnibus following the FAIR (Findable, Accessible, Interoperable and Reusable) Data Principles.

**Availability:** IRIS-EDA is freely available at http://bmbl.sdstate.edu/IRIS/.

**Supplementary information:** Supplementary data are available at *Bioinformatics* online.

## 1 Introduction

The development of high-throughput technologies, such as RNA-Seq, has created vast amounts of transcriptomic data that require efficient and scaled computational techniques (Wang, et al., 2009). One common investigation of RNA-Seq data is through analysis of estimated gene expression data. Analysis of the gene expression data is facilitated by computational experience in appropriately designing the methods and experiments and conducting the analysis processes using one of many computing languages. This creates an obstacle for users with limited computational experience who want to analyze their RNA-Seq studies, thus there is an increased need for easy-to-use interactive expression analyses and results visualization (Perkel, 2018).

While a wide variety of computational methods can be applied to expression data to determine particular qualities of the data on a sample or condition level (Abdi and Williams, 2010; Eisen, et al., 1998; Hartigan, 1972; Kruskal, 1964; Li, et al., 2009; Saelens, et al., 2018; Zhang, et al., 2016), differential gene expression (DGE) analysis is the most commonly used one. It allows researchers to identify differentially expressed genes (DEGs) across two or more conditions and can provide a meaningful way to attribute differences in gene expression levels to observed phenotypical and treatment differences. Many tools have been developed and optimized, such as: DESeq (Anders and Huber, 2012), DESeq2 (Love, et al., 2014), edgeR (Robinson, et al., 2010), limma (Ritchie, et al., 2015), Cuffdiff (Trapnell, et al., 2012), Cuffdiff2 (Trapnell, et al., 2013), sleuth (Pimentel, et al., 2017), and many others. While there have been substantial efforts in DGE analysis and visualization of DGE results (Ge, 2017; Goff, et al., 2013; Harshbarger, et al., 2017; McDermaid, et al., 2018; Nelson, et al., 2017; Nueda, et al., 2017; Powell, 2015; Younesy, et al., 2015), numerous pitfalls and bottlenecks persist, including experimental design implementation difficulties, a need for comprehensive integrated discovery-driven analyses and DGE tools, and the lack of functionalities and interactivity related to visualizing the analysis results.

To address these bottlenecks, we have created IRIS-EDA, which is an **I**nteractive **R**NA-Seq **I**nterpretation **S**ystem for **E**xpression **D**ata **A**nalysis. It provides a user-friendly interactive platform to analyze gene expression data comprehensively and to generate interactive summary visualizations readily. In contrast to other analysis platforms, IRIS-EDA provides the user with a more comprehensive and multi-level analysis platform. IRIS-EDA outperforms other tools in several critical areas related to efficiency and versatile applicability: 1) Single-cell and bulk RNA-Seq analysis capabilities, 2) GEO submission compatibility, 3) six useful discovery-driven and DGE analyses, 4) experimental design approaches through three integrated tools for DGE analysis, and 5) seven interactive visualizations (Figure 1A).

Specifically, IRIS-EDA provides comprehensive RNA-Seq data processing and analysis in a seamless workflow. This investigative approach uses expression quality control and discovery-driven analyses integrated with DGE analysis through one of the three most common R-based DGE tools (*Supplementary Materials S1*), *DESeq2, edgeR*, and *limma*, all of which have demonstrated capacities for expression data analysis (Sahraeian, et al., 2017). It provides users with a choice of intuitive experimental design options, as well as, the option to upload a custom design matrix in the DGE analysis. IRIS-EDA includes numerous interactive visualizations for each analysis type, enabling users to gain an immediate global view of their data and results or download as a high-resolution static image for publications. For the first time, this tool implements a framework based on the FAIR Data Principles (Wilkinson, et al., 2016) to assist users with the submission of their data and results to NCBI’s Gene Expression Omnibus (GEO).

IRIS-EDA is open source and freely available for use at http://bmbl.sdstate.edu/IRIS/ or can be loaded through R on a local computer. More information regarding implementation and accessibility can be found in *Supplementary Materials S2*.

**Figure 1:**
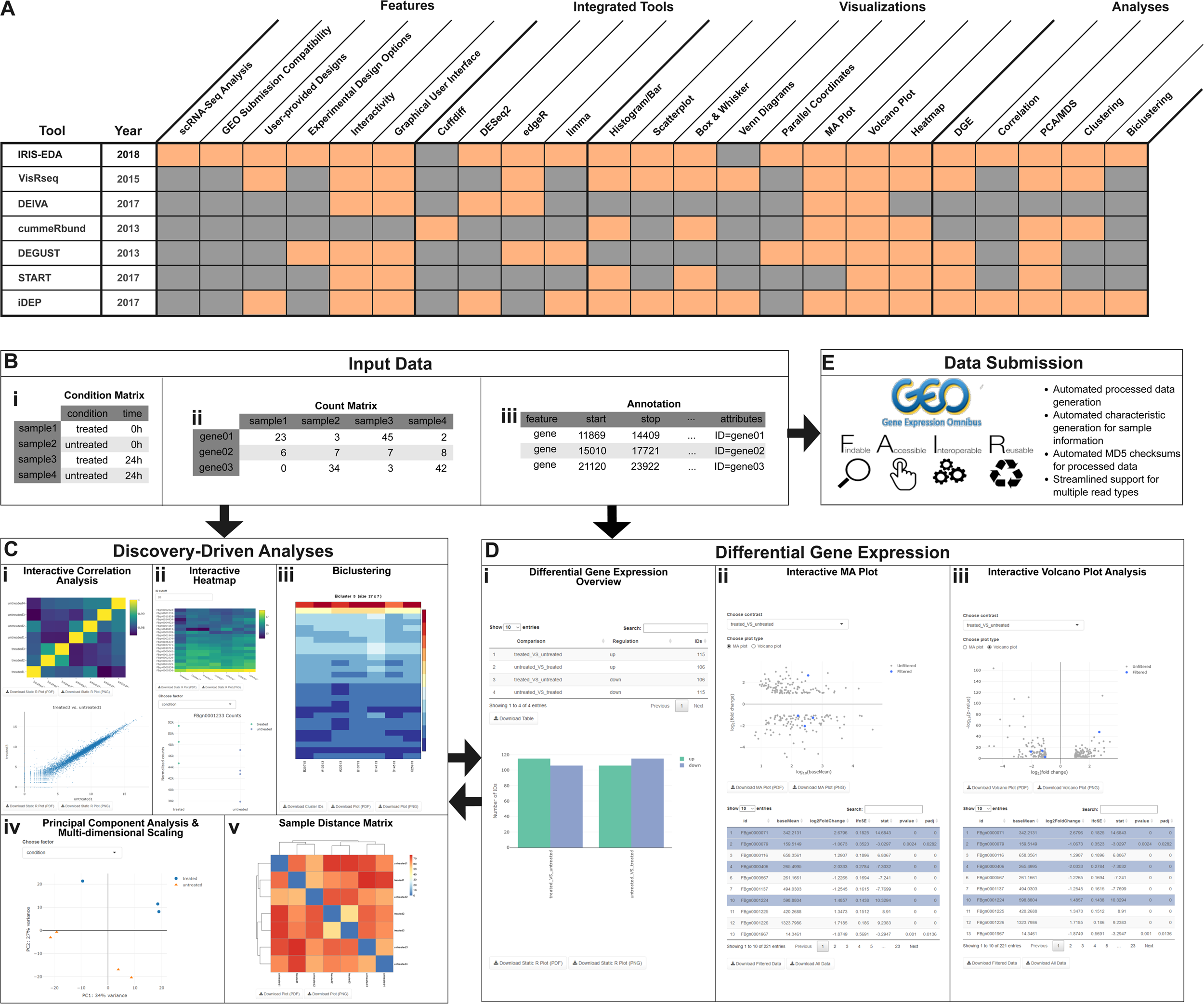
IRIS-EDA integrated functions. (A) Comparison of IRIS-EDA and six other DGE analyses and visualization tools; (B) Required Input Data for IRIS-EDA: (i) Condition Matrix indicating factor levels for each sample, (ii) Count Matrix consisting of gene expression values for each sample, with corresponding sample IDs matching those in the condition matrix, and (iii) the appropriate annotation file, which is required when using scRNA-Seq data; (C) Discovery-driven Analyses conducted by IRIS-EDA utilizing the Condition and Count matrices, including (i) Interactive Correlation Analysis with pairwise expression scatterplot, (ii) Interactive heatmap with parallel coordinate plot, (iii) Biclustering, (iv) Principal Component Analysis and Multi-dimensional Scaling, and (v) Sample Distance Matrix with clustering dendrogram; (D) Integrated Differential Gene Expression analysis with visualizations: (i) Differential Gene Expression Overview with table and bar charts corresponding to up- and down-regulated gene counts, (ii) Interactive MA Plot with DGE results table, and (iii) Interactive Volcano Plot with DGE results table; and (E) Data submission compatibility to Gene Expression Omnibus following the FAIR guiding principles.

## 2 Methods

### Bulk and Single-Cell RNA-Seq Analysis

IRIS-EDA was designed to provide a comprehensive platform for gene expression data analysis, which includes applicable analysis of both bulk and single-cell RNA-Seq data. Single-cell RNA-Seq (scRNA-Seq) data analysis is a growing area of study within RNA-Seq analyses and can provide unique insights into genetic occurrences within single cell types (McCarthy, et al., 2017; Soneson and Robinson, 2018). The methods used for traditional DGE analysis have demonstrated applicability to scRNA-Seq DGE analysis, under certain conditions (Soneson and Robinson, 2018). Thus, while designed for bulk RNA-Seq data analysis, IRIS-EDA can also facilitate discovery-driven and DGE analysis for scRNA-Seq data with few modifications. Namely, analysis of single-cell data can be appropriately carried out by using a stringent filter cutoff based on a default setting of transcripts per million (TPM) > 1, especially when combined with either edgeR or limma, which have both been shown to have high performance on scRNA-Seq data (Soneson and Robinson, 2018). More details regarding the analysis of scRNA-Seq data can be found in *Supplementary Materials S3*.

### Required Inputs

IRIS-EDA requires two or three user-provided input files, depending on the type of data used (Figure 1B and *Supplementary Materials S4*): (1) a gene expression estimation matrix (EEM, also referred to as sample count data), (2) a condition matrix with factor levels corresponding to the provided samples in the EEM, and (3) a gene length matrix indicating the base-pair length of each gene to be used for filtering of scRNA-Seq data only. When uploading their data, users will select whether they are uploading bulk or single-cell RNA-Seq gene expression data. If using scRNA-Seq data, the additional requirement for gene length matrix will be shown. Also, default parameterizations for optimized analysis for single-cell data will be populated throughout the server. Methods to obtain gene lengths from GFF/GTF/GFF3 annotation files can be found in *Supplementary Materials S3.2*.

Once users have uploaded required data, IRIS-EDA provides two distinct analysis approaches. First, users can explore their data through a comprehensive discovery-driven analysis approach. This method provides users with tools and analyses for exploratory analysis of their expression data. Second, users can perform differential gene expression (DGE) analysis on their submitted data. In this method, users can determine which genes are differentially expressed using one of the three integrated DGE tools and can visualize the results through interactive visualizations. Whether users choose to first analyze their expression data using the discovery-driven analyses or through DGE analysis, they can continue to investigate their data with the other approach as well, in order to provide a comprehensive view of their RNA-Seq expression data.

### Gene Expression Data Quality Control

After data upload, the two or three input files are first analyzed by IRIS-EDA quality control (*Supplementary Materials S5*). Input data quality is evaluated using boxplots and histograms of the read count distributions. The purpose of the quality control process is to enable exploration of the submitted data and to verify that there are no unexpected or unexplainable abnormalities in the data, such as low total read counts or individual samples displaying strange distribution behavior. Once users have established proper data quality, they can proceed to the investigative analyses provided in IRIS-EDA.

### Discovery-Driven Analyses

IRIS-EDA discovery-driven analyses (Figure 1C) are various tools and algorithms designed to provide an investigative approach of expression data, especially for the situation where users do not have a strong direction or hypothesis for their data analysis procedures. These algorithms assist users in analyzing and visualizing their EEM input information and discovering trends in their data that may provide additional hypotheses for downstream analyses. In particular, discovery-driven analyses can help users define a specific hypothesis within their RNA-Seq study, which can assist in development of experimental design methods for DGE analysis. Discovery-driven analyses processes that can be performed in IRIS-EDA include: sample correlation analysis and pairwise expression scatterplots (Figure 1Ci), expression heatmaps (Figure 1Cii), biclustering (Figure 1Ciii, QUBIC (Zhang, et al., 2016)), principal component analysis and multidimensional scaling (Figure 1Civ), and sample distance matrix with clustering (Figure 1Cv). The figures generated through the discovery-driven analysis feature of IRIS-EDA are provided in an interactive manner, allowing users to select specific samples or pairwise comparisons to further evaluate. One such example is with the sample correlation analysis and pairwise scatterplots shown in Figure 1Ci. Users can choose one cell of the sample correlation matrix corresponding to a comparison between two samples. This will display the pairwise scatterplot for that specific comparison. The user can then scroll over the scatterplot and display the gene ID for an indicated data point. Full details for each discovery-driven analysis and their respective interactivity can be found in the *Supplementary Materials S6*.

### Differential Gene Expression Analysis

After submitting data, users can move onto the DGE phase of IRIS-EDA (*Supplementary Materials S7*). This analysis is performed using any one of the three provided tools: *DESeq2* (Anders and Huber, 2012), *edgeR* (Robinson, et al., 2010), and *limma* (Ritchie, et al., 2015). The default tool is *DESeq2,* based on independent evidence supporting its performance (Sahraeian, et al., 2017) and our RNA-Seq analysis experience, but users can also select one of the other two tools based on their own preference. There are other high-performing commonly-used DGE tools available; however, their compatibility with IRIS-EDA excludes their use in IRIS-EDA. For example, tools that do not utilize sample count data, e.g., *Sleuth,* (Pimentel, et al., 2017) or are not R-based, e.g., *Cuffdiff* (Trapnell, et al., 2012), are not included due to compatibility issues.

In addition to the DGE tool, the experimental design can also be specified by the user. The designs provided in IRIS-EDA include two-group comparisons for analysis of selected pairwise comparisons, multiple factorial comparisons, classic interaction design, additive models for pairing or blocking of data, main effect testing (testing time-series data) and blocked main effect testing. Additionally, IRIS-EDA provides a method for users to specify their own experimental design, for the instances when the user needs a design not already included in IRIS-EDA. Each of these methods has unique parameters to specify by the user, typically including which factors are intended for analysis and which specific comparisons are required. After analyzing the data, IRIS-EDA provides an overview displaying the number of up- and down-regulated IDs for each indicated comparison, along with a histogram displaying this information (Figure 1Di). The results table is also available through IRIS-EDA, along with interactive MA (Figure 1Dii) and Volcano plots (Figure 1Diii).

Similar to the figures generated in the Discovery-Driven Analysis section of IRIS-EDA, the plots in the DGE section are also highly interactive. Discovery-Driven Analysis features allows users to gain more specific information from their plots, including highlighting individual or regions of data points on the plot. These features highlight the corresponding row of the DGE results table, showing users gene information identifying them as outliers or falling within a certain region. Conversely, users can select specific gene IDs from the results table, resulting in the highlighting of that gene ID’s or set of gene IDs’ data points on the corresponding plot. This feature can be used to easily determine the relative location of specific genes or gene sets in the plot.

Results obtained from the DGE analysis section of IRIS-EDA are often not the end of the analysis procedures. Based on the information collected, users may choose to further investigate their expression data using additional analyses provided in the Discovery-Driven Analyses section, such as the clustering or biclustering. When DGE and Discovery-Driven analyses are combined, the analyses provide a more comprehensive data interpretation.

### IRIS-EDA Outputs

IRIS-EDA provides users with methods for extracting content based on discovery-driven and DGE analyses. All figures in the QC, Discovery-Driven Analysis, and DGE Analysis sections have the option for users to download as a static image in PDF or PNG format. Additionally, all tables in the DGE Analysis section are downloadable as CSV files, with the final results table being downloaded in its entirety or filtered based on user-provided or default-adjusted p-value and log fold-change cutoffs. As part of the biclustering analysis, users can also download a list of gene IDs contained within the specified cluster.

### GEO Submission and FAIR Data Principles compatibility

Many users are eventually interested in submitting their RNA-Seq data to a public repository for accessibility, but this process can be tedious and troublesome. NCBI’s GEO database has specific requirements related to the data, results, and accompanying metadata file. To assist users in their preparation of documents for GEO submission, IRIS-EDA offers an optional GEO page. In following with the standard of set forth by the FAIR Data Principles (Wilkinson, et al., 2016), this page asks users to provide a limited amount of information that will be used, along with the previously provided condition matrix information, to populate the metadata file required for GEO submission. This populated metadata file will then be available for download with reformatted processed data files extracted from the EEM. These two pieces of information can later be submitted with the original raw FASTQ-formatted RNA-Seq data to the GEO submission page. More detailed information regarding the usage of the GEO capabilities of IRIS-EDA can be found in *Supplementary Materials S8*.

## 3 Conclusions

IRIS-EDA is a platform developed for comprehensive expression data analysis, visualization and interpretation of both bulk and single-cell RNA-Seq data. It is designed to address current bottlenecks and issues in existing expression analysis and DGE analysis packages. This interactive tool implements numerous features including EEM quality control, discovery-driven analyses, and DGE analysis utilizing the most commonly used R-based DGE tools in a user-friendly, comprehensive platform. It is noteworthy that IRIS-EDA provides advanced experimental design options in an intuitive format, while also allowing users to provide their own design matrix to facilitate efficient DGE analysis for a broad spectrum of users. Each analysis section within IRIS-EDA provides relevant information in a highly-interactive visual format. To further facilitate compatibility with the FAIR Data Principles, IRIS-EDA also provides a framework that will greatly assist users in formatting their results and metadata for GEO submission. It is our belief that this tool will support users of all computational experience levels and with all DGE requirements.

