## Supplementary Materials for "IRIS-EDA: An integrated RNA-Seq interpretation system for gene expression data analysis" (from another version of this preprint: version 1, 2018-03-16)

### IRIS-DGE Supplementary Material

[1]South Dakota State University
[2]Department of Biology and Microbiology
[3]Bioinformatics and Mathematical Biosciences Lab
[4]Department of Mathematics and Statistics
[5]Population Health Group, Sanford Research
[6]Department of Internal Medicine, Sanford School of Medicine
[7]Department of Agronomy, Horticulture, and Plant Science
[8]BioSNTR

**03/16/2018**

**Abstract**

Next-Generation Sequencing has made available much more large-scale genomic data. Studies with RNA-sequencing (RNA-seq) data typically involve generation of genetic expression profiles that can be further analyzed, many times involving differential gene expression (DGE). This process enables a comparison across samples of two or more factor levels. A recurring issue with DGE analyses is the complicated nature of the comparisons to be made, in which a variety of factor combinations, pairwise comparisons, and main or blocked main effects need to be tested. Here we present a tool called IRIS-DGE, which is a server-based DGE tool developed using Shiny providing a straightforward, user-friendly platform for performing comprehensive DGE analysis. This tool provides crucial analyses that help design hypotheses and to determine key genomic features. IRIS-DGE integrates the three most commonly used R-based DGE tools to determine differentially expressed genes (DEGs) and includes numerous methods for performing preliminary analysis on user-provided genetic expression information. Additionally, this tool integrates a variety of visualizations, in a highly interactive manner, for improved interpretation of preliminary and DGE analyses.

### S1: Citation information

**Table 1:** A comparative overview of citation counts for DGE tools and DGE servers. DGE analytical tools (*Tool*) are compared based on the following criteria: Current number of citations (*Citations*), percentage of total citations from the analytical tools presented (*Citation %*), year the analytical tool was published (*Year*), approximate citations per year based on data accrued through 2017 (*Citations/Year*), and if the analytical tool has an R-based application (*R-based*).

| Tool | Citations | Citation % | Year | Citations/Year (through 2017) | R-based? |
|---|---|---|---|---|---|
| edgeR (Robinson et al. 2010) | 7093 | 32.39700375 | 2010 | 1013.285714 | Yes |
| Cuffdiff (Trapnell et al. 2012) | 4533 | 20.70430255 | 2012 | 906.6 | No |
| Cuffdiff2 (Trapnell et al. 2013) | 1508 | 6.887731799 | 2013 | 377 | No |
| DESeq2 (Love et al. 2014) | 4249 | 19.40714351 | 2014 | 1416.333333 | Yes |
| limma (Ritchie et al. 2015) | 2399 | 10.95733991 | 2015 | 1199.5 | Yes |
| DEGseq (Wang et al. 2009) | 1237 | 5.649949758 | 2009 | 154.625 | Yes |
| baySeq (Hardcastle et al. 2010) | 562 | 2.56691331 | 2010 | 80.28571429 | Yes |
| SAMseq (Li et al. 2013) | 275 | 1.256051886 | 2013 | 68.75 | Yes |
| NOIseq (Tarazona et al. 2012) | 38 | 0.173563533 | 2012 | 7.6 | Yes |

### S2: Accessibility

IRIS-DGE can be freely accessed directly through this link or through R using the following code in the proceeding sections.

#### S2.1: Install CRAN packages

IRIS-DGE requires several packages to operate. To get these pacakages, the following commands can be entereed into an R terminal to check if you already have the necessary packages. If not, the following code will install any missing packages:

```
# CRAN
packages <- c(
    "crosstalk", "dplyr", "DT", "gtools", "plotly", "shiny", "plyr",
    "shinyBS", "shinycssloaders", "shinythemes", "tibble", "tidyr",
    "Rcpp", "Hmisc", "ggplot2", "geneplotter", "locfit", "GGally",
    "pheatmap", "reshape2", "backports", "digest", "fields", "psych"
)
np <- packages[!(packages %in% installed.packages()[, "Package"])]
if(length(np)) install.packages(np)
```

#### S2.2: Install Bioconductor packages

You will also need several Bioconductor packages. Similar to the prior section, the following code will check and install any missing Bioconductor packages into your R library:

```r
# Bioconductor
bioc.packages <- c("DESeq2", "edgeR", "limma", "QUBIC")
np <- packages[!(bioc.packages %in% installed.packages()[,"Package"])]
source("https://bioconductor.org/biocLite.R")
if (length(np)) biocLite(np)
```

#### S2.3: Run the Shiny application

Once you have installed all of the necessary packages, you can now run the application locally by entering the following code into an R terminal:

```r
shiny::runGitHub("iris", "btmonier")
```

Typically, the link will provide an easier route to using IRIS-DGE. In circumstances where internet connections will be limited (such as during travel), loading IRIS-DGE through R while internet is still available will allow users to utilize IRIS-DGE without an internet connection later on.

Installing the local application via GitHub will also let the user have access to more "developmental" versions of IRIS-DGE. Be warned though; using developmental versions of this application may be "cutting edge", however, this could potentially break various features in the application. Therefore, using the server (http://bmbl.sdstate.edu/IRIS/) will provide a more "stable", peer-reviewed release of the application.

### S3: A note about input data

IRIS-DGE requires two pieces of information for analysis. The first is an expression estimation matrix, also referred to as a count matrix, displaying the gene expression estimates for each sample. The format requires a CSV file with the row names to list the gen IDs and column names to list the sample IDs. The second required input is a condition matrix, wherein the factor levels for each sample are provided. This file requires a CSV format and row names to be the sample IDs matching the sample IDs from the expression estimation matrix and the column names to be the condition factors.

The data used for this tutorial are derived from 28 *Vitis vinifera* (grape) samples with three distinct factors (Rootstock, row, and block). This data can viewed as the "big" example data set found under the `Submit and QC` tab under `1. Submission Parameters`.

#### S3.1: Expression matrices

Typically, an expression matrix, also known as count data or a count matrix refers to data where every $i$-th row and $j$-th column refer to how many reads are assigned to gene (ID) $i$ in sample $j$. For example, if we have simplified count data for 4 samples and three genes, the `R` output will look something like this:

```
        sample1 sample2 sample3 sample4
gene001      23       3      45       2
```

```
gene002        6        7        7        8
gene003        0       34        3       42
```

**Note:** When loading count data into IRIS-DGE, make sure that the first column is your gene IDs and that sample names are short, concise, and avoid the use of mathematical operators (`+`, `-`, `/`, `*`, `^`, etc.) and spaces between words. If a space is necessary for legibility, please consider using an underscore (`_`)

#### S3.2: Condition matrices

Condition matrices, also known as metadata, details the design of your experiment. In this type of matrix, every $i$-th row and $j$-th column refer to factor levels assigned to sample $i$ and factor $j$. For example, if were to look at the samples given in the count data section, the metadata `R` output will look something like this:

```
        condition time
sample1   treated   0h
sample2 untreated   0h
sample3   treated  24h
sample4 untreated  24h
```

**Note[1]:** When loading metadata into IRIS-DGE, make sure that the first column is your sample names and that column names and treatment levels are short, concise, and avoid the use of mathematical operators (`+`, `-`, `/`, `*`, `^`, etc.) and spaces between words. If a space is necessary for legibility, please consider using an underscore (`_`)

**Note[2]:** Metadata can be expanded to fit the nature of your experiment (i.e. multiple factors can be added). The only thing that must remain consistent between these two matrices, is the sample information. Column names in count data **must** be the same as row names in the metadata.

### S4: Expedited analysis

#### S4.1: Overview

Expedited analysis is for users who want a quick and efficient method of producing DGE results using the default parameters and tools in IRIS-DGE.

#### S4.2: Submit and QC

1. Click on the `Submit and QC` tab:

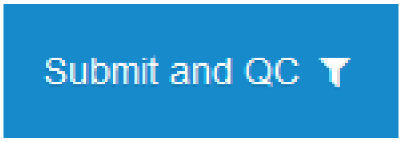

2. Load user data by selecting `Load my own data` under `1. Submission Parameters`. User data requires one count matrix and one condition matrix. **Note:** you must supply both forms of data:

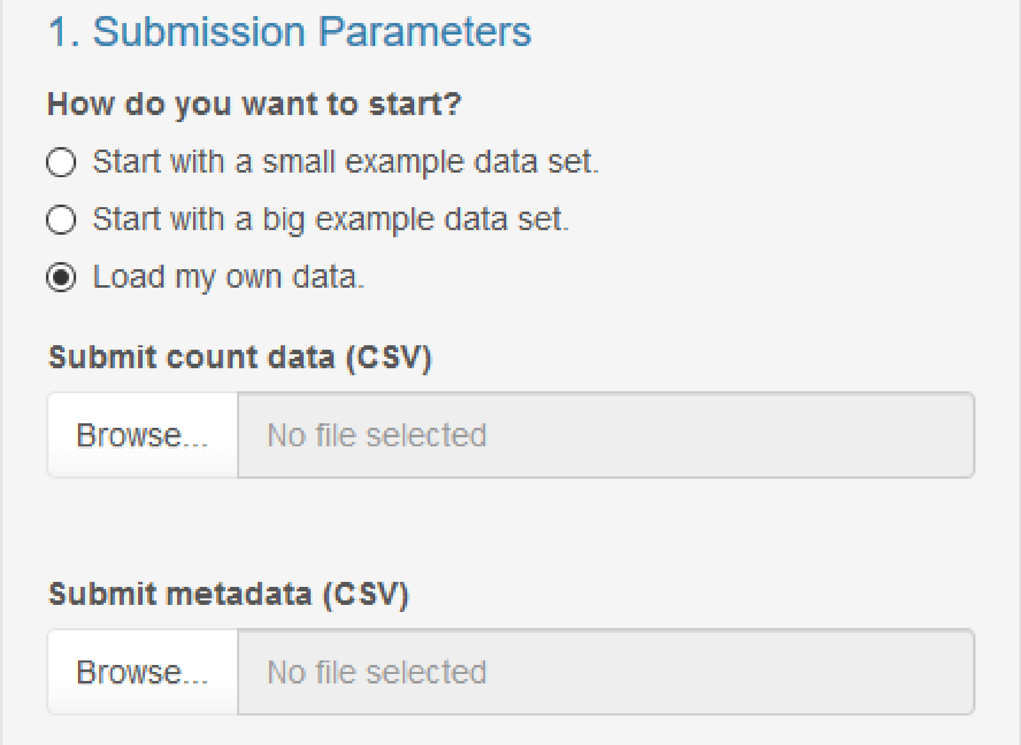

3. Click `Submit` to load the data under `3. Launch Overview`:

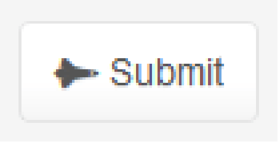

4. After submitting the data, proceed to the `DGE Analysis` tab at the top. **Note:** if a specific experimental design other than the basic two-group comparison is required, select it accordingly:

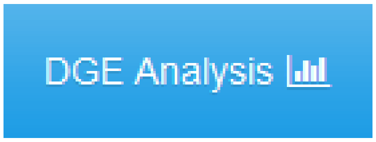

5. Select the factor of interest and check the boxes for the comparisons of interest under `3. Experimental Parameters`. **Note[1]:** you must choose at least one comparison for whatever experimental design you choose under `1. Experimental Setup`! **Note[2]:** If you do not want to use `DESeq2`, feel free to change this method to either `edgeR` or `limma-voom` under `2. DGE Paramters`. Also, feel free to change adjusted $p$-values and minimum fold change if these defaults do not fit your analysis:

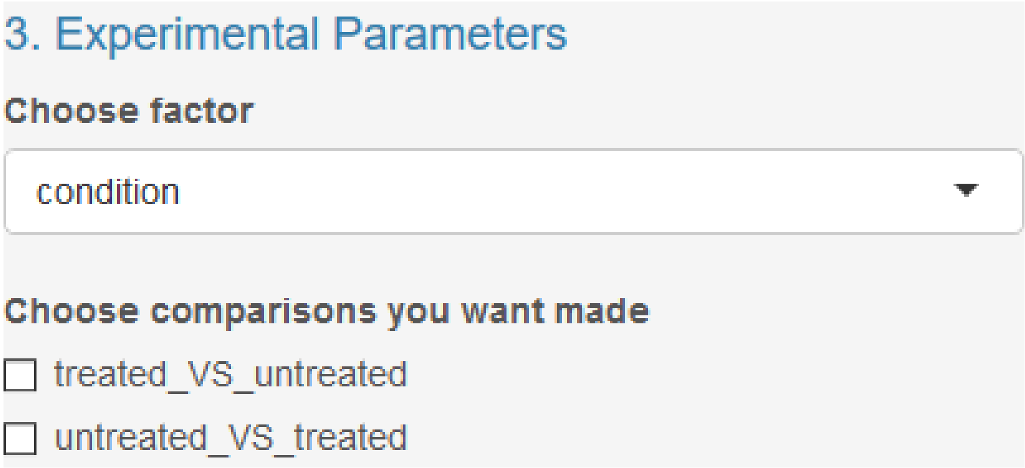

5. Submit the parameters to perform DGE analysis under `4. Launch Analysis`:

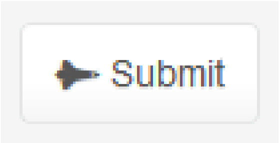

6. Select the newly formed `Plots` subtab on the main page to view the DGE results table towards the bottom:

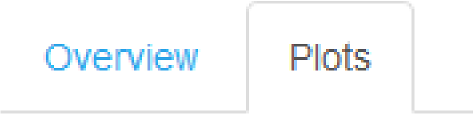

#### S4.3: Data export

1. To export results data from the `DGE Analysis` tab, click on the `Download All Data` button at the bottom of the page to download the results file:

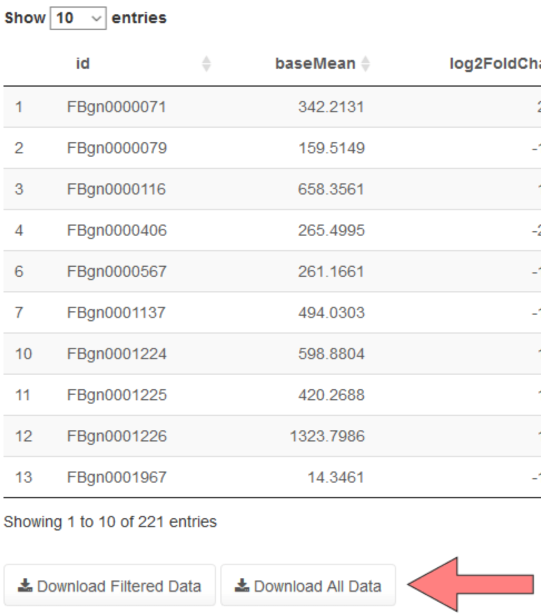

| | id | baseMean | log2FoldChange | lfcSE | stat | pvalue | padj |
|---|---|---|---|---|---|---|---|
| 1 | FBgn0000071 | 342.2131 | 2.6796 | 0.1825 | 14.6843 | 0 | 0 |
| 2 | FBgn0000079 | 159.5149 | -1.0673 | 0.3523 | -3.0297 | 0.0024 | 0.0282 |
| 3 | FBgn0000116 | 658.3561 | 1.2907 | 0.1896 | 6.8067 | 0 | 0 |
| 4 | FBgn0000406 | 265.4995 | -2.0333 | 0.2784 | -7.3032 | 0 | 0 |
| 6 | FBgn0000567 | 261.1661 | -1.2265 | 0.1694 | -7.241 | 0 | 0 |
| 7 | FBgn0001137 | 494.0303 | -1.2545 | 0.1615 | -7.7699 | 0 | 0 |
| 10 | FBgn0001224 | 598.8804 | 1.4857 | 0.1438 | 10.3294 | 0 | 0 |
| 11 | FBgn0001225 | 420.2688 | 1.3473 | 0.1512 | 8.91 | 0 | 0 |
| 12 | FBgn0001226 | 1323.7986 | 1.7185 | 0.186 | 9.2383 | 0 | 0 |
| 13 | FBgn0001967 | 14.3461 | -1.8749 | 0.5691 | -3.2947 | 0.001 | 0.0136 |

### S5: In-depth analysis

#### S5.1: Overview

By choosing the in-depth analysis route, you will be given more information about your data including:

- Count data distribution;
- Total number of reads/sample;
- Sample correlation analysis;
- Biclustering;
- Principal components and multidimensional scaling;
- Identification of most variable IDs in data.

#### S5.2: Submit and QC

1. Click on the `Submit and QC` tab near the top-left corner of the application:

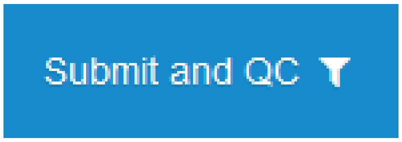

2. Under `1. Submission Paramters`, select either `Start with a small example data set`, `Start with a big example data set`, or `Load my own data` to upload user data. **Note:** User data requires one count matrix and one condition matrix:

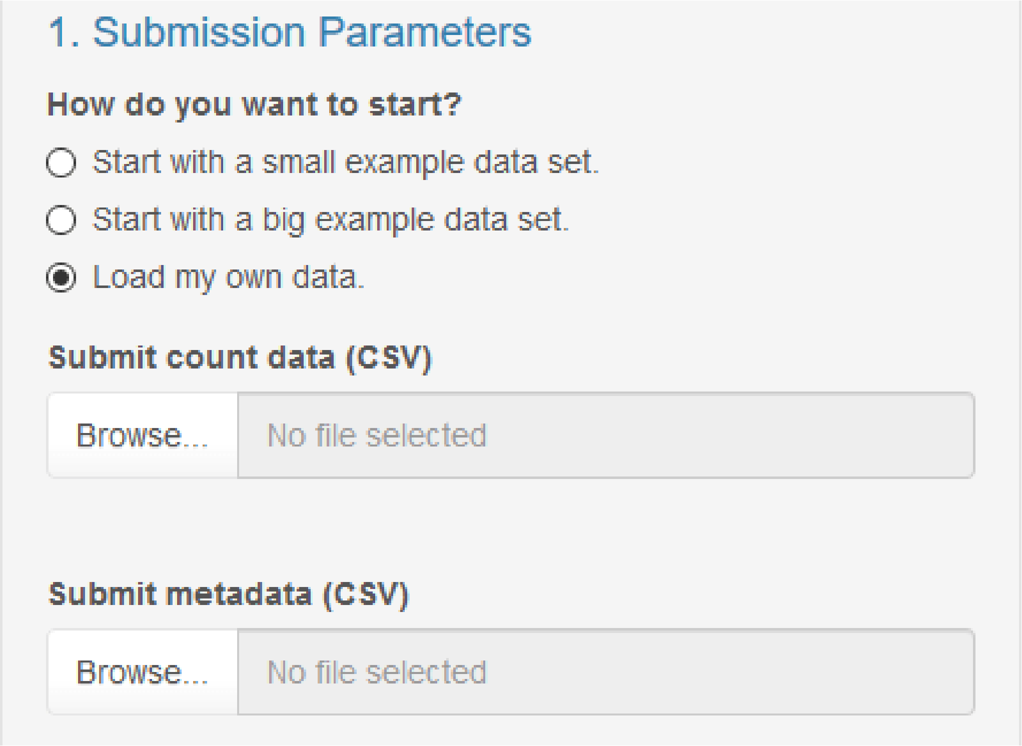

2. Under `2. Data Processing`, select a filter cutoff to simplify and expedite computations by eliminating any rows that have below specified expression totals. The default argument for our application is `10`:

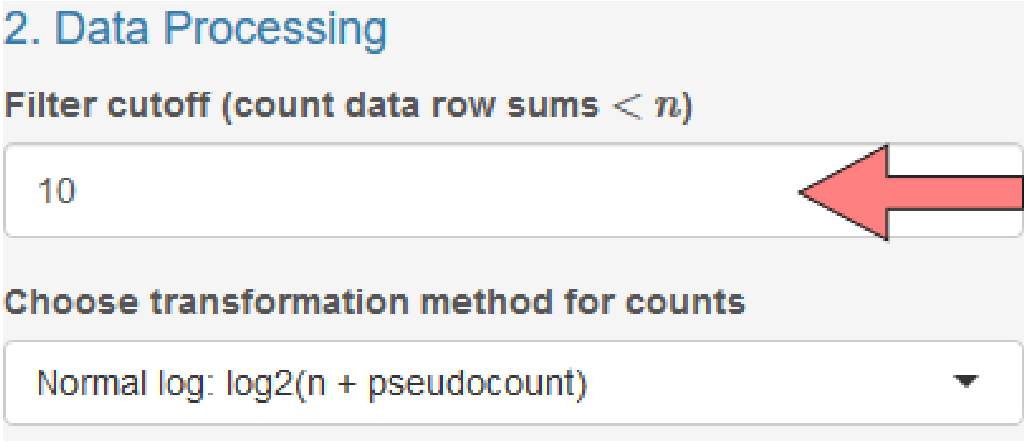

3. Right below the filter cutoff parameter, select a transformation method for the count data for displaying the expression estimate. **Note:** For more information about any of these topics, take a look at the *FAQ* section at the bottom of this document:

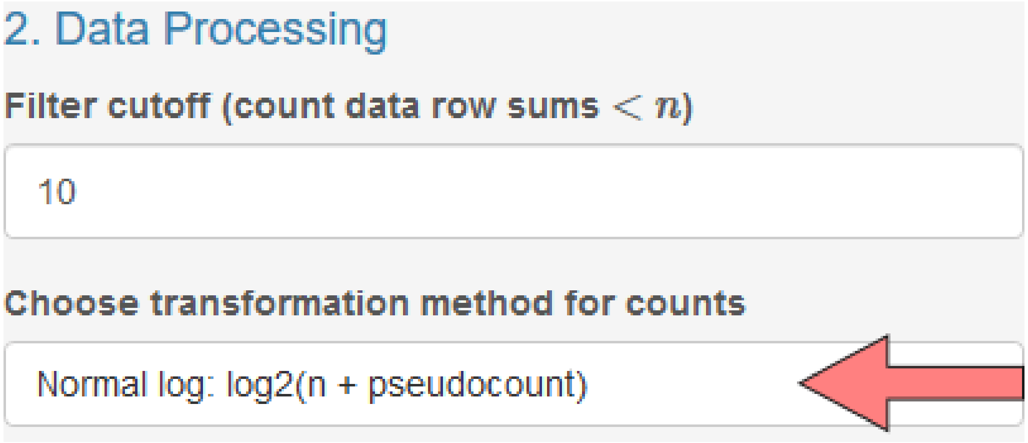

4. Click `Submit` to load the data under `3. Launch Overview`:

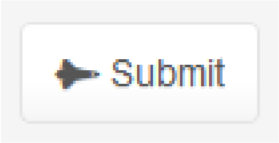

5. After you click `Submit`, the main page will now be populated with several pieces of information in two sub-tabs, `File Summary` and `Count Summary`.

#### S5.2.1: File Summary

`File Summary` provides a glimpse into the submitted data. This subtab includes three main components.

1. The `count data` section will detail the first and last five rows of the count data and also include the total number of IDs and samples:

##### Count data (first 5 rows and last 5 rows)

```
Your pre-processed data contains 42412 IDs and 28 samples. Showing the first 10 samples:

               A10115 B10515 B20715 C10915 D11315 D21515 E21915 F12115 G12515 G22715
4CL             16934  12829  18027  12877   6697  10380  12540  10025  16769  13998
ABP1              483    598    461    537    411    474    707    457    673    780
CALS10          10680  12684  10841  14071   7965   8374  14314  11046  12090  14358
CBL01             445    566    729    692    417    377    805    504    702    621
CBL02            1338   1514   1665   1734   1028   1137   1969   1356   1538   1739
...               ...    ...    ...    ...    ...    ...    ...    ...    ...    ...
XM_010663230.1     96     82    154    126    127    105    463     46    160    610
XM_010666890       54     73     43     75     57     59     67     54     78     90
XM_010667047      656    956    728    912    570    554   1091    720    826   1063
XM_010667194      894    978   1086   1252    738    730   1087    860    991   1100
vitvi15g00009    2032   2329   2933   2741    988   1443   2591   2258   2203   2367
```

2. The second portion includes an overview of the condition data this is simply an `R` console-based output of this submitted CSV file:

#### Sample metadata

| | Rootstock | Block |
|---|---|---|
| A10115 | OWN | A15 |
| B10515 | C3309 | B15 |
| B20715 | C3309 | B15 |
| C10915 | P1103 | C15 |
| D11315 | SO4 | D15 |
| D21515 | SO4 | D15 |
| E21915 | C3309 | E15 |
| F12115 | SO4 | F15 |
| [illegible] | [illegible] | [illegible] |

3. Finally, pre- and post-filtered gene ID counts gives a "before-and-after" count of the number of IDs that were filtered using the filter cutoff parameter under `2. Data Processing`:

#### Number of IDs (pre-filtration)

```
[1] 42412
```

#### Number of IDs (post-filtration)

```
[1] 27892
```

##### S5.2.2: Count Summary

`Count summary` provides three interactive, downloadable visualizations based on the file for each sample.

1. Box-and-Whisker Plot for transformed read counts by sample with interactivity showing the quartiles, maximum, and minimum count. *With the example data, it appears that the box-and-whisker plot for each sample is similar to the other samples. If one sample had a plot varying greatly from the others, it would indicate some required investigation into that specific sample in terms of the number of raw reads provided and proportion of reads aligned.*

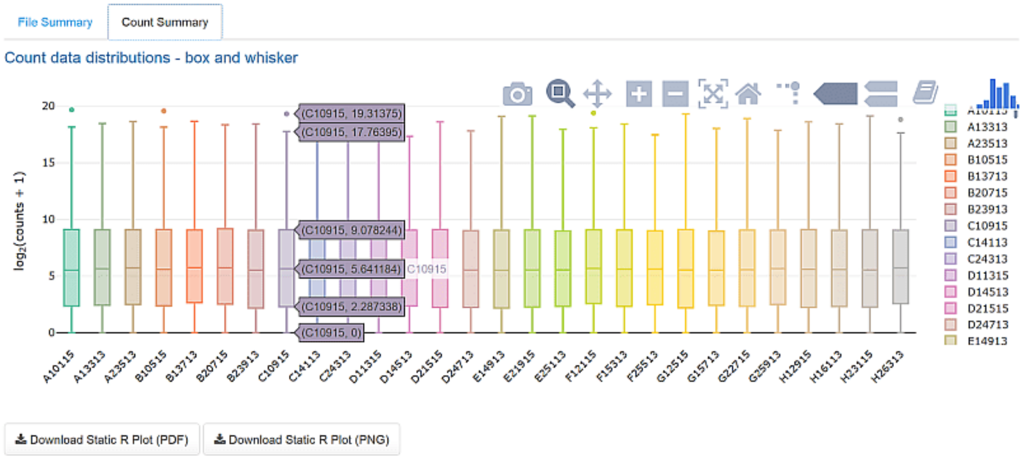

2. Count Data Distribution plot showing the frequency of transformed count data by sample with interactivity displaying the value and frequency. Additionally, double-clicking a sample ID in the legend isolates just that sample's histogram. Additional sample IDs can be select for more specific comparisons also. *With the example data, the histograms appear similar for each sample, indicating no significant derivation. Similar to the box-and-whisker plots, a single sample varying greatly from other samples may indicate a required investigation into the raw read counts or read alignment statistics.*

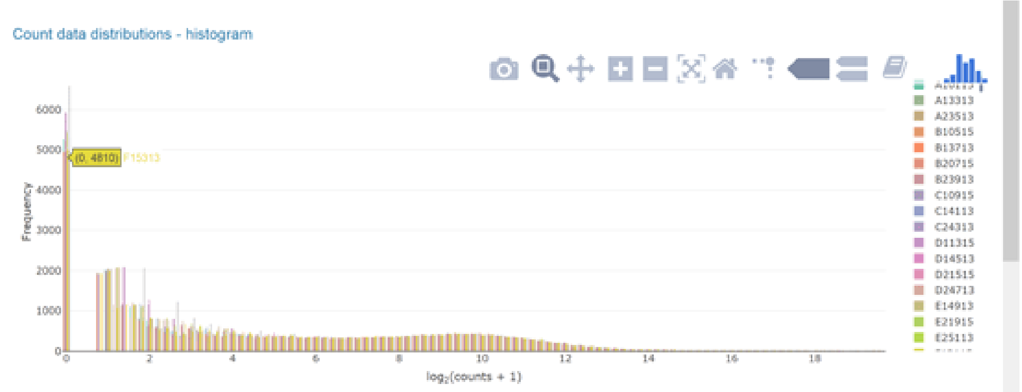

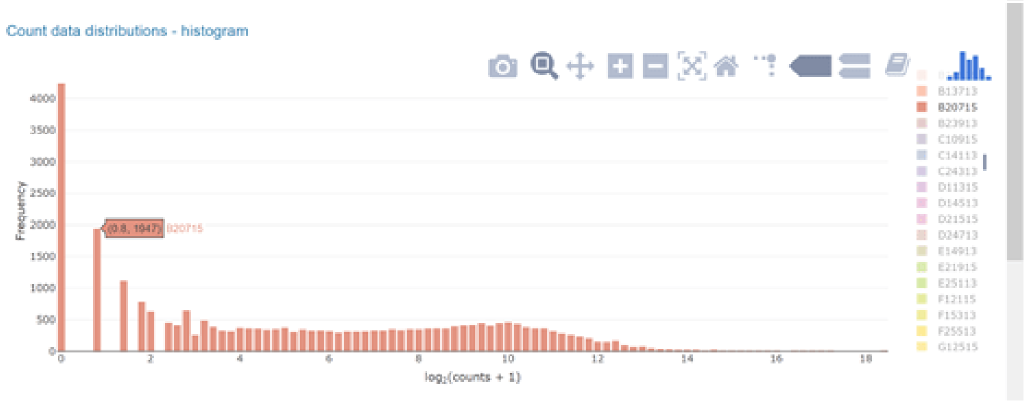

3. Total Reads displays a histogram of total reads by sample with interactivity for displaying actual total read counts for each sample. Double-clicking on a sample ID in the legend isolates that sample's read count histogram and allows for selecting of specific adjacent

sample IDs for comparison. Total reads counts for individual samples that vary greatly from the other total read counts may indicate some issues with data preparation (sequencing) or read alignment. *Here, sample H26313 has a much lower total reads count than the other samples. This may be reflected in further comparative analyses.*

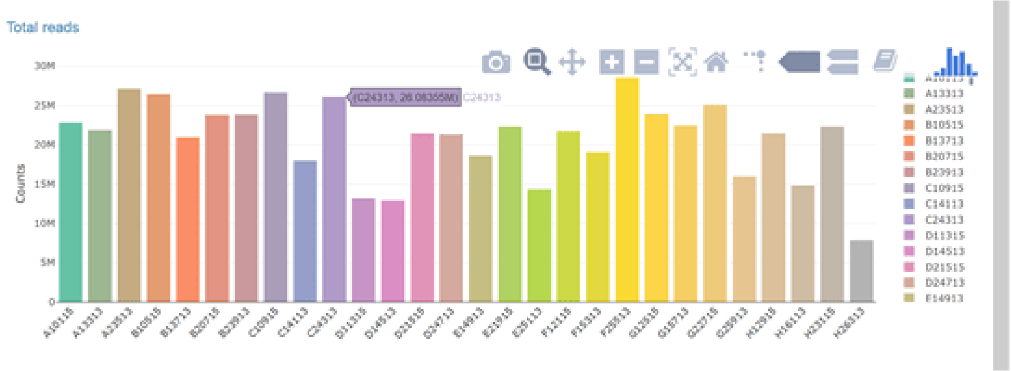

#### S5.3: Preliminary Analysis

1. After examining your results on the `Submit and QC` tab, you may proceed to the `Preliminary Analysis` tab:

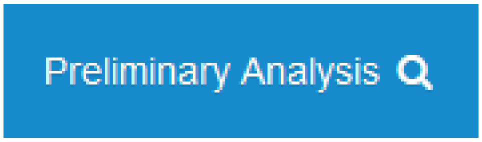

2. The `Preliminary Analysis` tab will be populated with several subtabs, similar to the `Submit and QC` tab. The first subtab you will see is the `Correlation` tab. This tab provides correlation analysis of the expression matrix by sample:

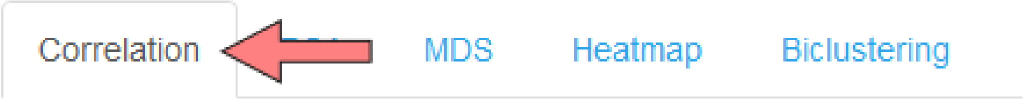

3. Under the `Correlation` subtab you will see several visualizations. `Interactive Correlation Analysis` displays a heatmap of the correlation between samples with interactivity showing the actual correlation between the two intersecting samples. *The example data shows most sample-sample correlations of 0.95 or larger, indicating relatively high correlation. The darker cells here signify less similar samples, which may yield more interesting differential expression results. This graph can indicate comparisons of interest in future analyses. In most cases, the high number of gene IDs with similar or identical expression estimates will cause correlations to be large, even for dissimilar genetic expression profiles:*

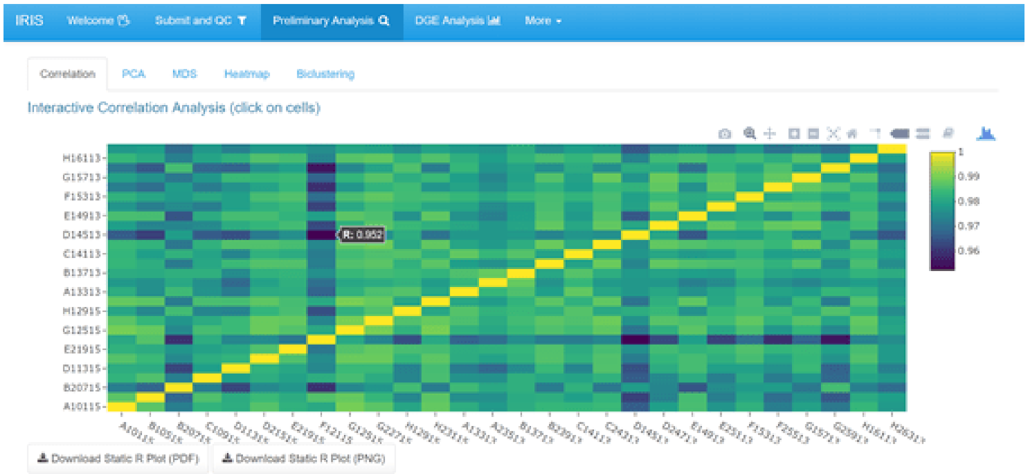

4. Clicking on a cell will provided a scatterplot of the two intersecting samples with each data point representing one gene ID. A scatterplot with points falling more closely along the diagonal indicates samples with more similar genetic expression profiles. *This scatterplot shows a clear trend of data points (gene IDs) falling along or close to the diagonal. That means these two samples have very similar genetic expression profiles. Data points that fall far from the diagonal line represent genes with dissimilar expression levels between the select samples:*

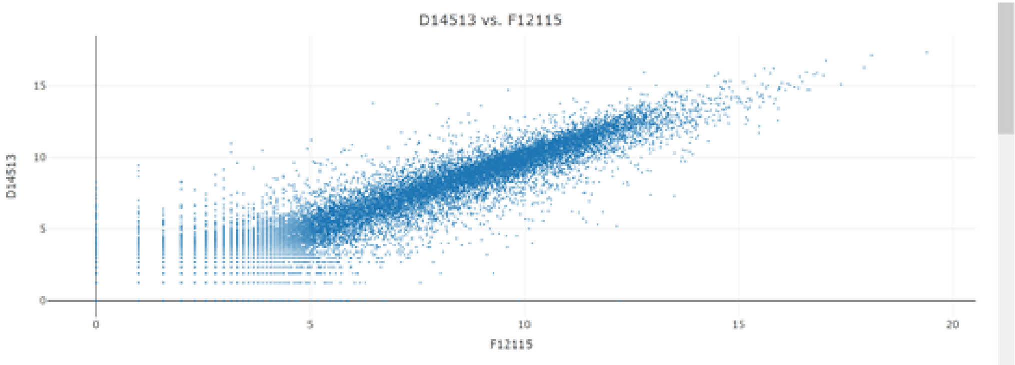

5. The Sample Distance Matrix provides a heatmap of the Euclidean distance between the gene expression vectors for each sample pair. The larger distance (darker red color) indicates samples with the most dissimilar genetic expression profiles. This matrix also includes a clustering of the samples based on the vectorized expression profiles. *With the example data, two distinct clusters can be observed through the first branching of the dendrogram. Additionally, as with the correlation heatmap, specific cells with a darker color indicate a more dissimilar pair of samples based on genetic expression:*

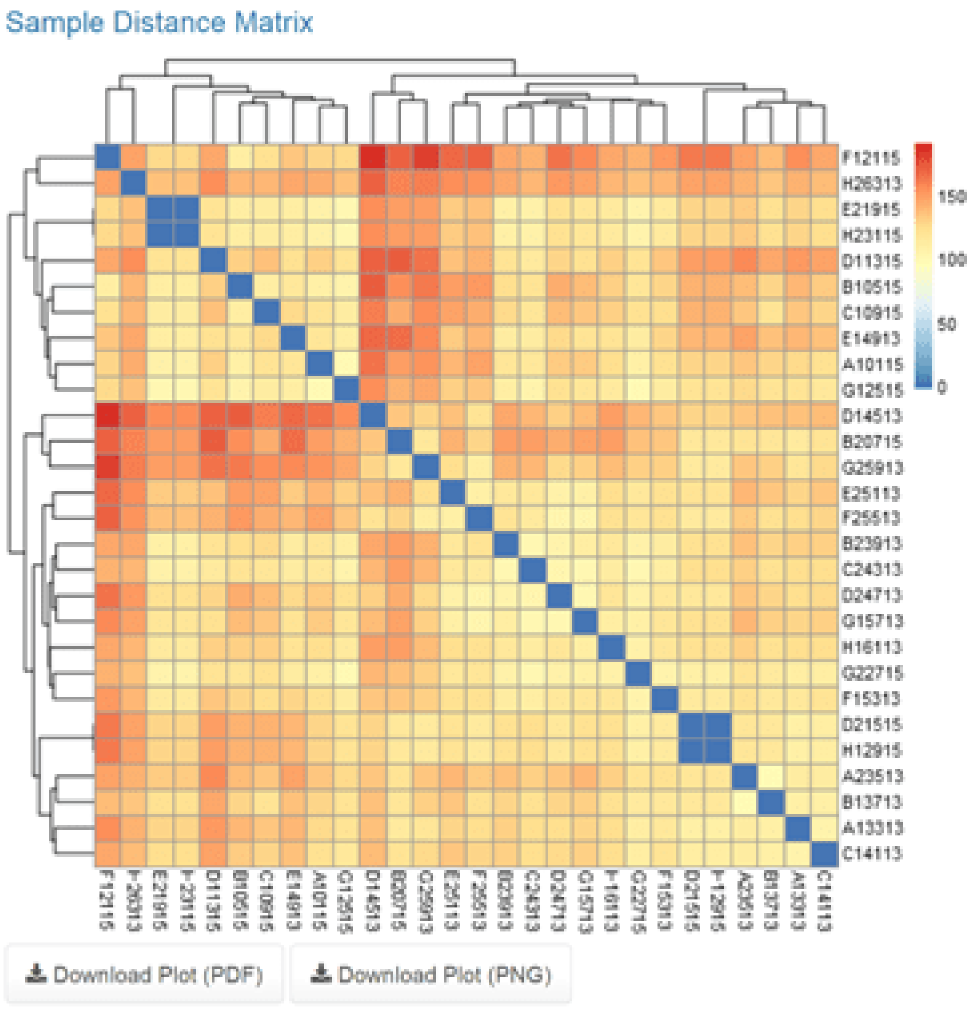

6. The next visualization will be under the `PCA` subtab. This subtab provides Principal Component Analysis (PCA) for the expression estimation matrix:

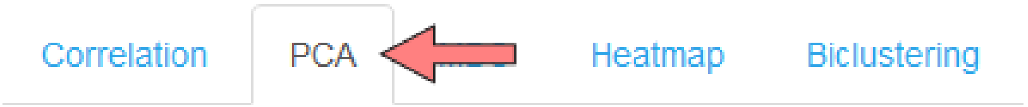

7. This analysis has the option of selecting a factor of interest. *With the example data, selecting "Rootstock" as the factor of interest provides a visualization of the first two components for each sample. In this application, PCA is a linear transformation of the gene expression levels, with the first component representing the transformed dimension with the most variability, and each subsequent component decreasing in variability. This analysis has the potential to isolate samples based on expression levels. Here, there does not appear to be any specific rootstock that separates from the others. If there were, it could help develop directions for further analysis. The axis labels indicate the first principal component accounts for 37% of the variance between samples, whereas the second principal component accounts for 7%:*

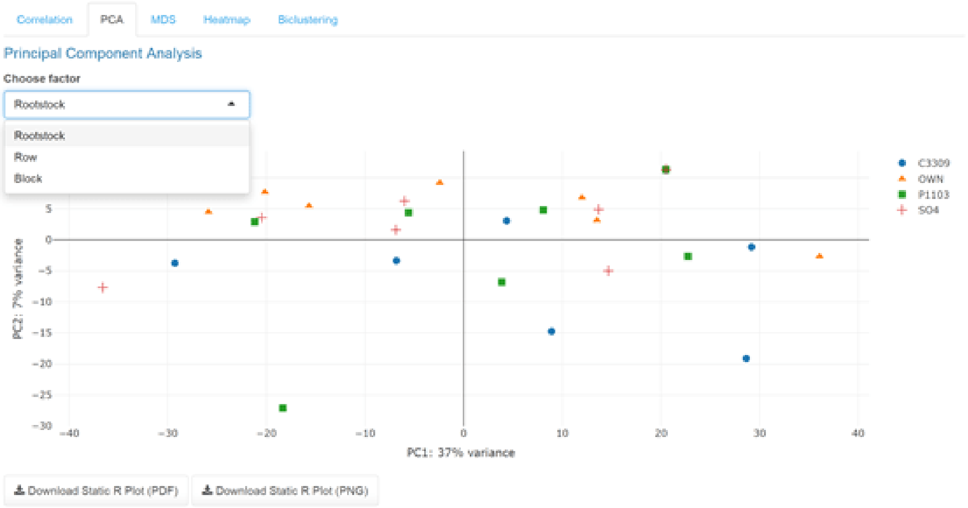

8. The next visualization will be under the `MDS` subtab. This subtab provides multi dimensional scaling for the expression estimation matrix. This is similar to PCA except is develops components through non-linear transformations of the gene expressions:

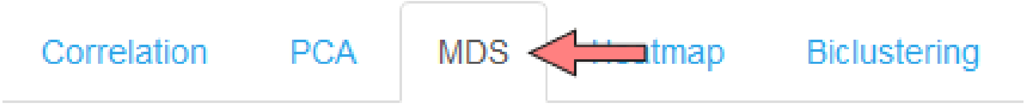

9. *Looking back at our sample data, we observe similar results to that of the PCA results, with similar potential interpretations if any sample or groups of samples were to differentiate from the others:*

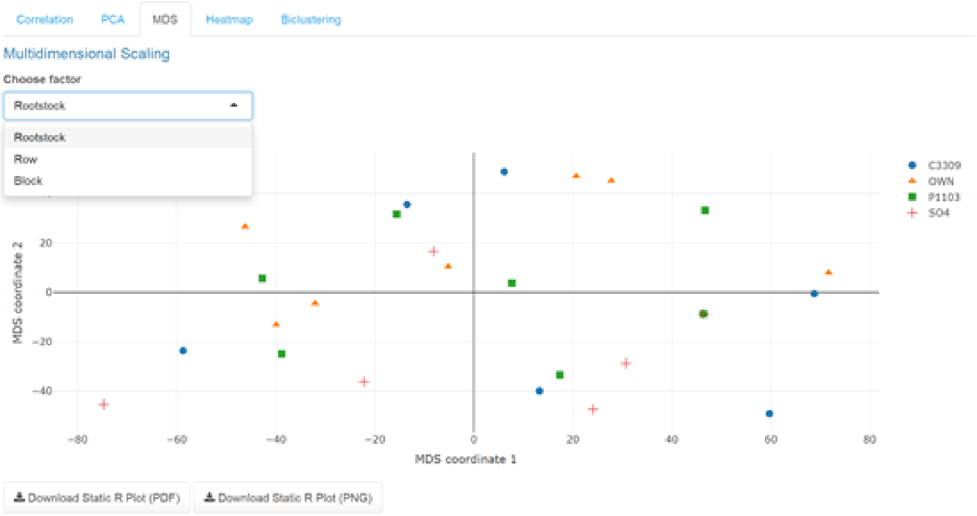

10. The next visualization will be under the `Heatmap` subtab. This subtab provides an interactive heatmap with rows representing the gene IDs and columns representing sample IDs:

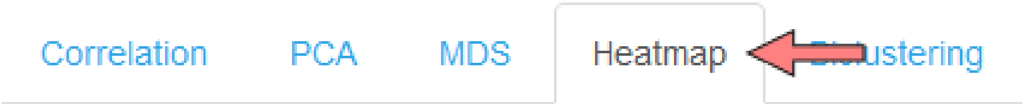

11. This heatmap requires an ID cutoff to select the indicated number of gene IDs with the highest mean expression values for display. Selecting a cell displays a plot showing the total read counts for that specific gene ID by the selected factor. *With the example data, the 20 most variable gene IDs are displayed. The yellow color indicates gene IDs with a higher expression level for that sample, and the darker blue color represents a low expression level for that sample. Selecting ID: rna25007 shows the read counts for that ID by rootstock factor. This shows the "OWN" rootstock seems to have a higher expression level for that ID, with the exception of one sample:*

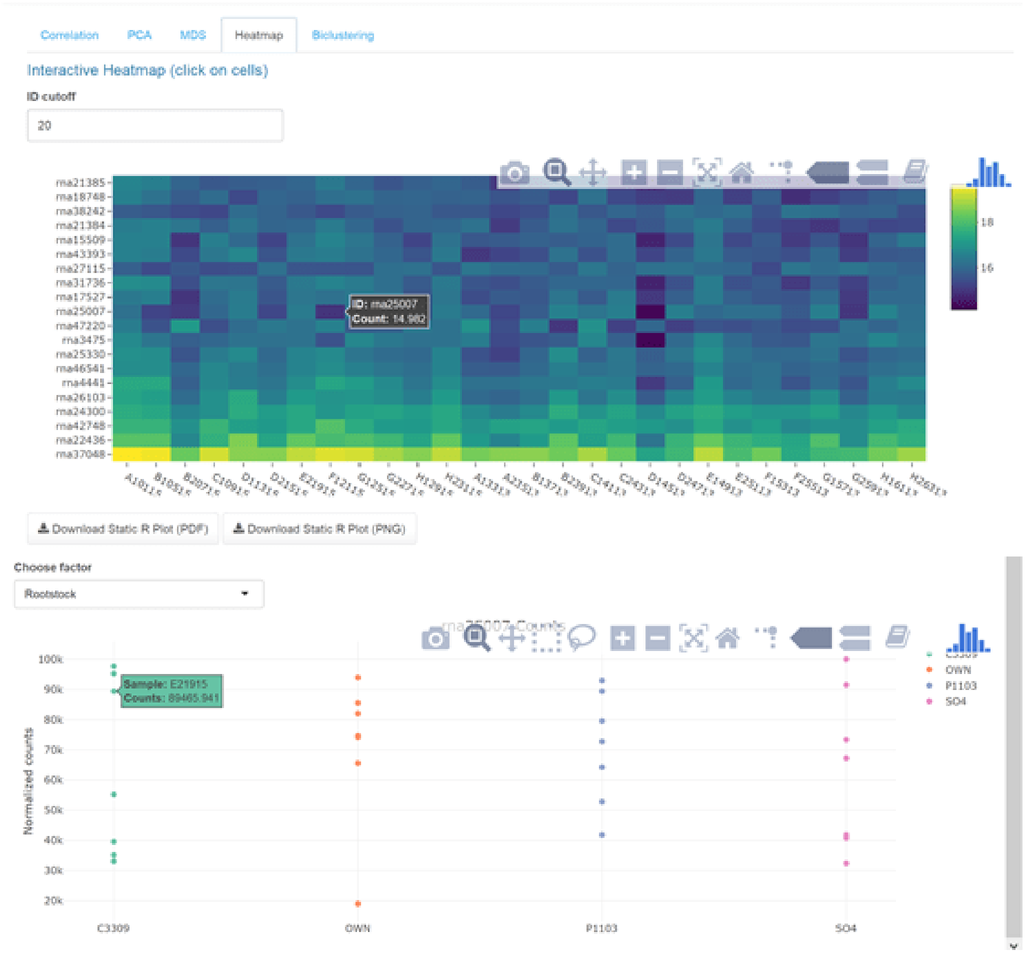

12. On the next subtab (1), `Biclustering` performs a biclustering analysis using one of the selected biclustering tools (3) with a maximum bicluster size of the indicated cutoff value (2):

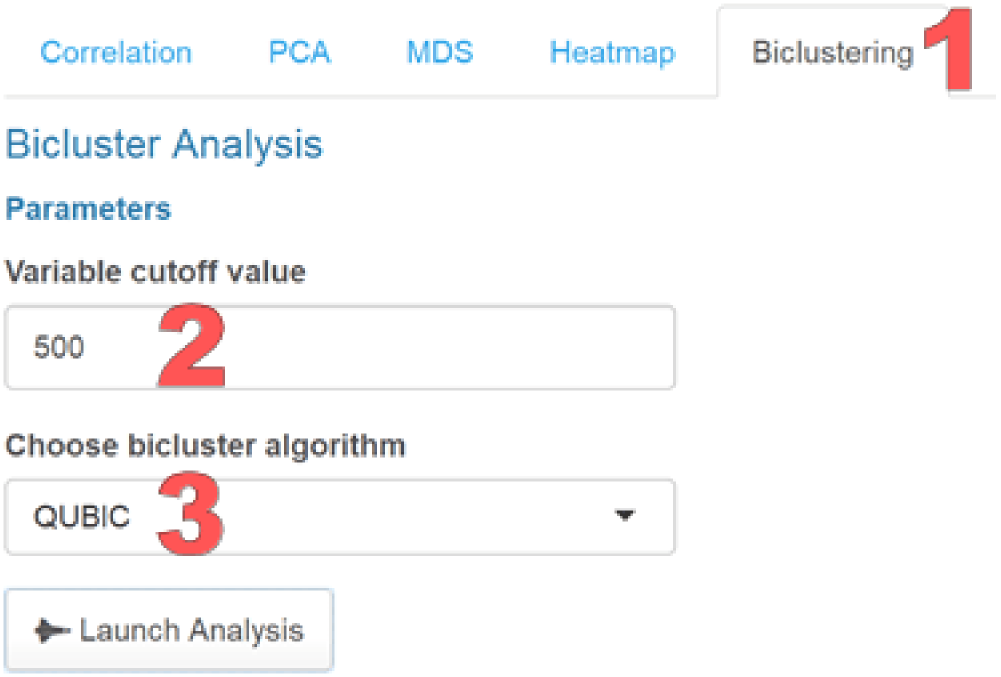

13. Launching the analysis results in display of the first bicluster. Alternative clusters can be selected from the dropdown menu and the IDs and plot for each bicluster can be downloaded using the button below the visualization. *With the sample data, the biclusters can help select the samples under which certain gene IDs are similarly expressed. Since gene expression levels can vary greatly over all samples and conditions, a biclustering approach can isolate similar expression patterns on a level where traditional clustering may miss. The first cluster for the example data shows that samples B20715, D21515, and H12915 are expressed similarly under the isolated gene IDs. Interpretations can be made similarly for each subsequent bicluster:*

##### Clustering Overview

Choose cluster

1

This algorithm found 93 IDs amongst 3 samples in cluster 1.

##### Heatmap analysis of cluster 1

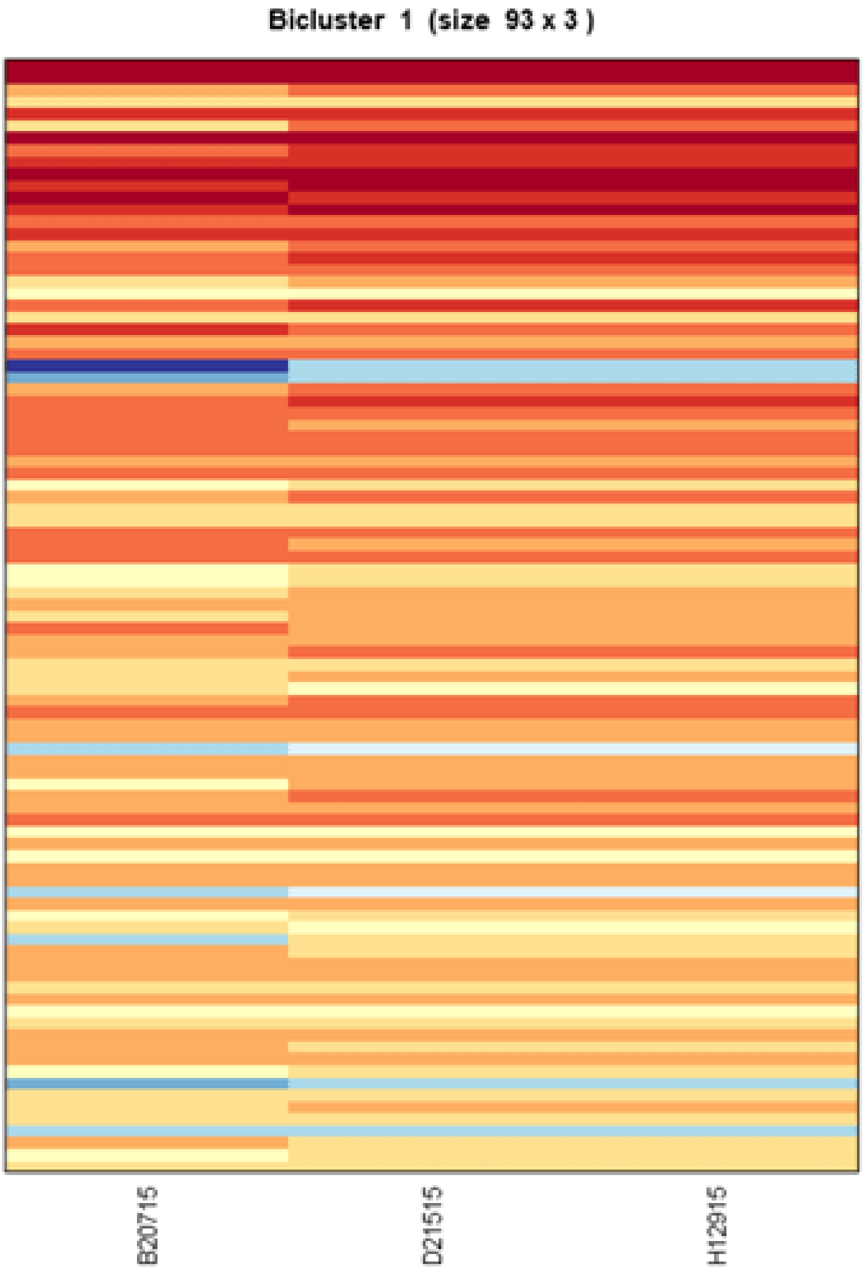

#### S5.4: DGE Analysis

1. After uploading the user data or selecting the example data, users can go directly to the `DGE Analysis` tab, preceding the `Preliminary Analysis` functions:

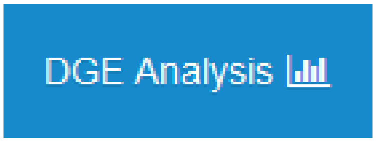

2. Once you are on this tab, you will be greeted with several options. `1. Experimental Setup` will allow users to select an experimental design for their DGE analysis. Options are Two-group comparison, Multiple factor comparisons, Classical interaction design, Additive models (paired or blocking designs), Main effects, Main effects with grouping factors, or Custom design:

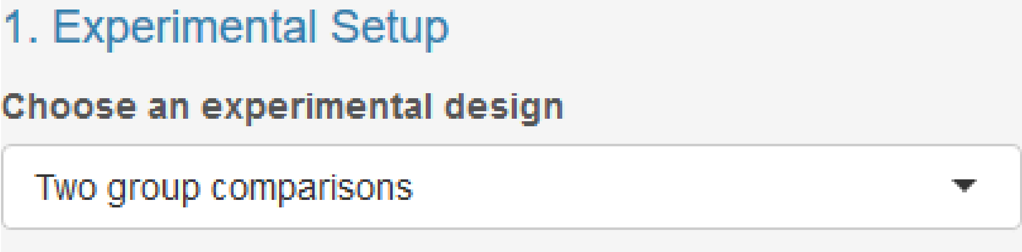

##### S5.4.1: An overview of experimental designs

1. The `Two-group comparisons` options is the traditional approach for DGE and compares two factors levels for the selected factor type. *With the example data, selecting "Two group comparisons" for the experimental design and "Rootstock" for the factor allows for specific pairwise comparisons of Rootstock factor levels. Here, we can select specific comparisons of interest from the permutations of all pairwise comparisons. Selecting all comparison options will provide inverse duplications, so specific selections may be needed. Below, all unique pairwise combinations are selected. The linear model is also displayed for users interested in the model used for comparisons:*

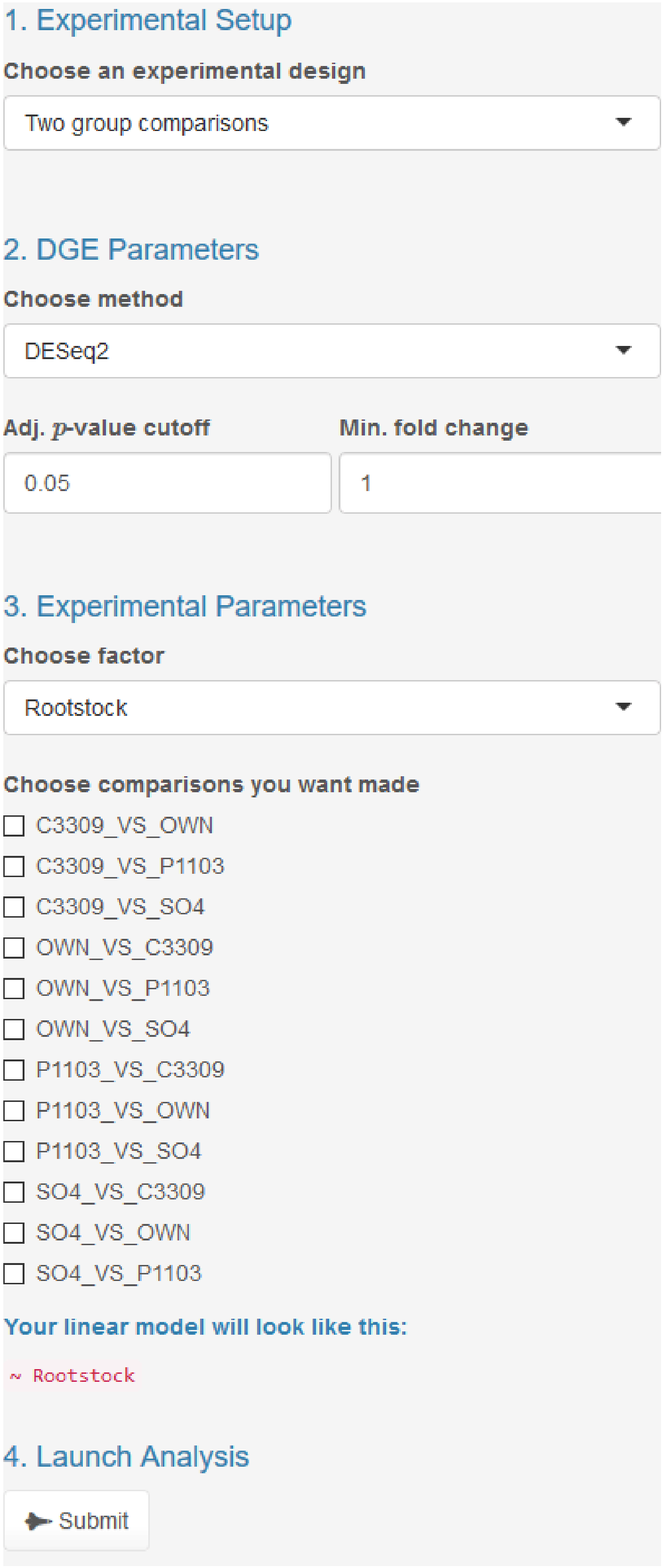
1. Experimental Setup
Choose an experimental design
Two group comparisons
2. DGE Parameters
Choose method
DESeq2
Adj. p-value cutoff
Min. fold change
0.05
1
3. Experimental Parameters
Choose factor
Rootstock
Choose comparisons you want made
C3309_VS_OWN
C3309_VS_P1103
C3309_VS_SO4
OWN_VS_C3309
OWN_VS_P1103
OWN_VS_SO4
P1103_VS_C3309
P1103_VS_OWN
P1103_VS_SO4
SO4_VS_C3309
SO4_VS_OWN
SO4_VS_P1103
Your linear model will look like this:
~ Rootstock
4. Launch Analysis
Submit

2. The `Multiple factor comparisons` design has users select two factor levels and performs all crosswise comparisons for the two chosen factor levels. *With the example data, the Multiple factor comparisons design with Rootstock and Block selected as the two factors provides optional comparisons for each rootstock separated by block. In this situation, as with the other designs, the user selects which comparisons are of interest. Selecting C3309_B_VS_C3309_E allows for a comparisons of gene expression levels for the same rootstock in two different blocks. This provides insight into the locations and possibly time (due to time requirements for sampling) for this specific rootstock:*

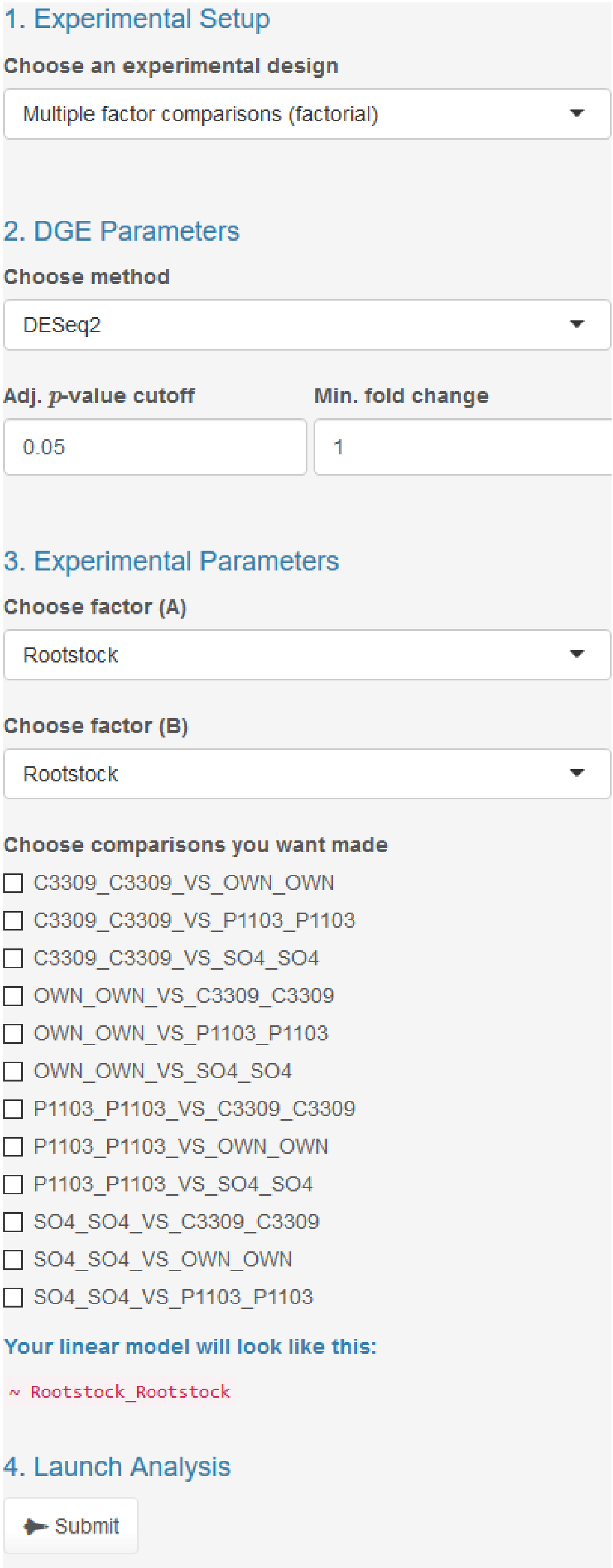
1. Experimental Setup
Choose an experimental design
Multiple factor comparisons (factorial)
2. DGE Parameters
Choose method
DESeq2
Adj. p-value cutoff
0.05
Min. fold change
1
3. Experimental Parameters
Choose factor (A)
Rootstock
Choose factor (B)
Rootstock
Choose comparisons you want made
C3309_C3309_VS_OWN_OWN
C3309_C3309_VS_P1103_P1103
C3309_C3309_VS_SO4_SO4
OWN_OWN_VS_C3309_C3309
OWN_OWN_VS_P1103_P1103
OWN_OWN_VS_SO4_SO4
P1103_P1103_VS_C3309_C3309
P1103_P1103_VS_OWN_OWN
P1103_P1103_VS_SO4_SO4
SO4_SO4_VS_C3309_C3309
SO4_SO4_VS_OWN_OWN
SO4_SO4_VS_P1103_P1103
Your linear model will look like this:
~ Rootstock_Rootstock
4. Launch Analysis
Submit

3. The `Classical interaction design` allows the user to select two factor levels and one reference level for each of the selected factors. *With the experimental data, the Classic interaction design allows for selection of two factors and a reference for each. In this case, the Rootstock and Row are the chosen factors and OWN and 15 are the selected reference levels for comparison. The contrast levels selected upon submission will provide DEGs with respect to these two levels:*

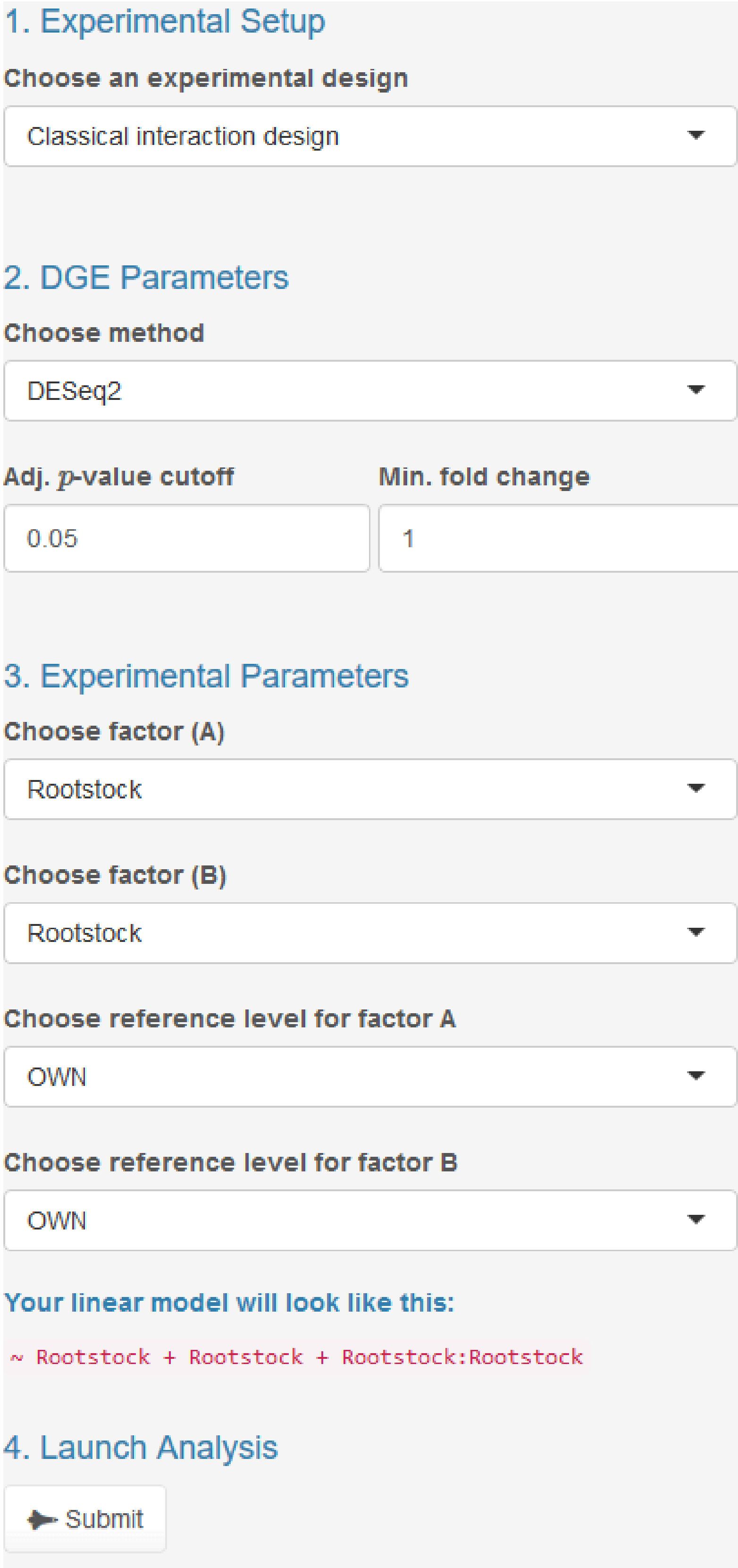
1. Experimental Setup
Choose an experimental design
Classical interaction design
2. DGE Parameters
Choose method
DESeq2
Adj. p-value cutoff
0.05
Min. fold change
1
3. Experimental Parameters
Choose factor (A)
Rootstock
Choose factor (B)
Rootstock
Choose reference level for factor A
OWN
Choose reference level for factor B
OWN
Your linear model will look like this:
~ Rootstock + Rootstock + Rootstock:Rootstock
4. Launch Analysis
Submit

4. The `Additive models` design is useful when samples are paired or blocked into distinct groupings. This format requires selected factors, one for the pairing or blocking factor and the other for the treatment factor. Additionally, a reference level for each selected factor is required. *In the example data, the Rootstock factor can be considered for grouping and the treatment factor is Block. OWN is selected as the reference level for the blocking factor and the A15 Block is selected for the treatment reference level:*

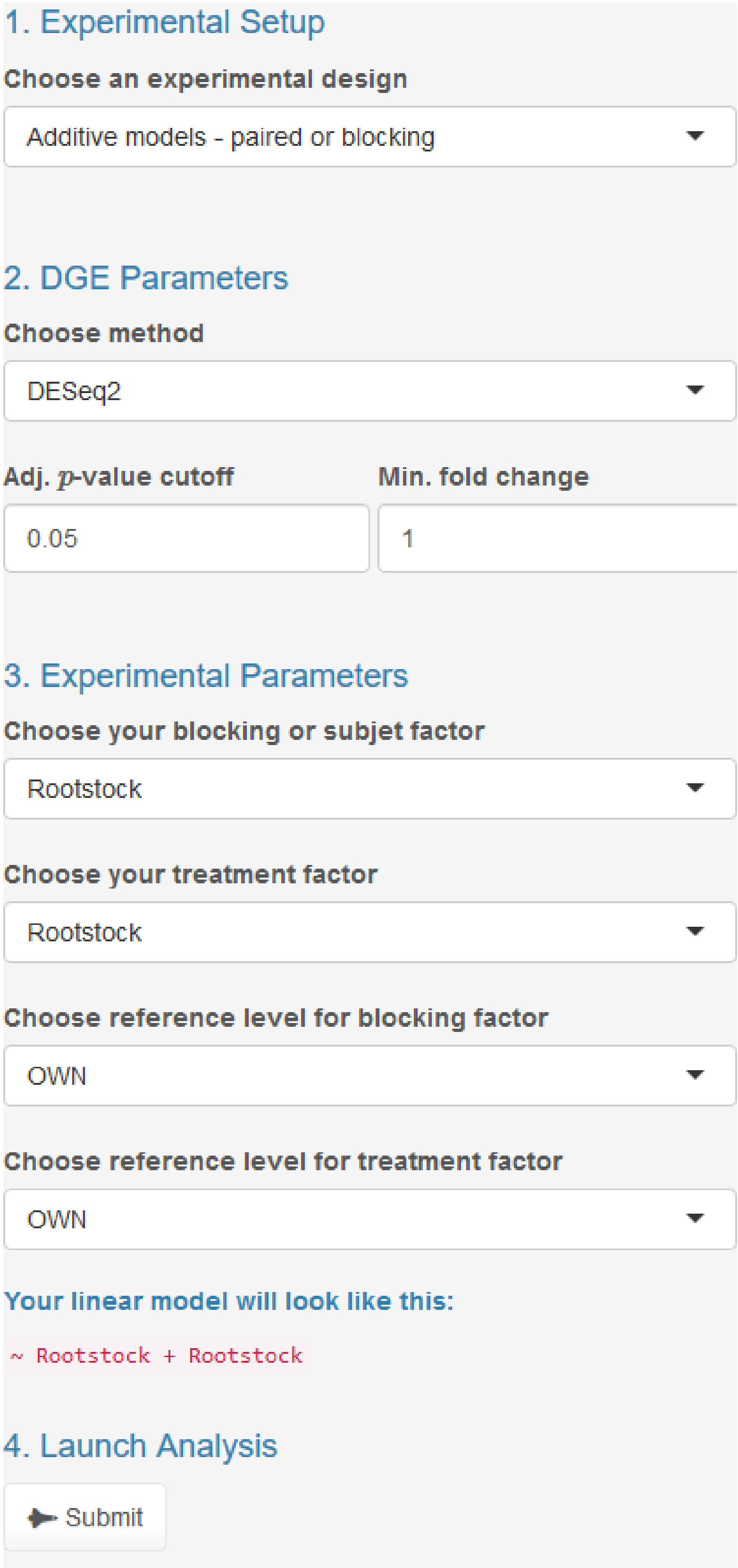
1. Experimental Setup
Choose an experimental design
Additive models - paired or blocking
2. DGE Parameters
Choose method
DESeq2
Adj. p-value cutoff
Min. fold change
0.05
1
3. Experimental Parameters
Choose your blocking or subjet factor
Rootstock
Choose your treatment factor
Rootstock
Choose reference level for blocking factor
OWN
Choose reference level for treatment factor
OWN
Your linear model will look like this:
~ Rootstock + Rootstock
4. Launch Analysis
Submit

5. `Main Effects` experimental design allows for testing the significance of a factor across multiple factor levels. In this situation, any significant deviation from an intercept would result in a significantly differential expressed gene. Using this approach is most useful when users want to test the significance of a factor that has more than two levels. This design requires indication of which factor to test as the main effect. Additionally, users must specify a factor reference level for the lfc and for corresponding visualizations. The $p$-values, adjusted $p$-values, and related differentially expressed genes are not affected by the selected reference level. *In the example data, the Rootstock factor is selected, which will test whether Rootstock has a significant impact on the genetic expression across all Rootstock factor levels. Selection of the C3309 Rootstock factor level as a reference will provide lfc values and visualizations relative to this level, while $p$-values and adjusted $p$-values will be calculated with respect to the main effect of the chosen factor:*

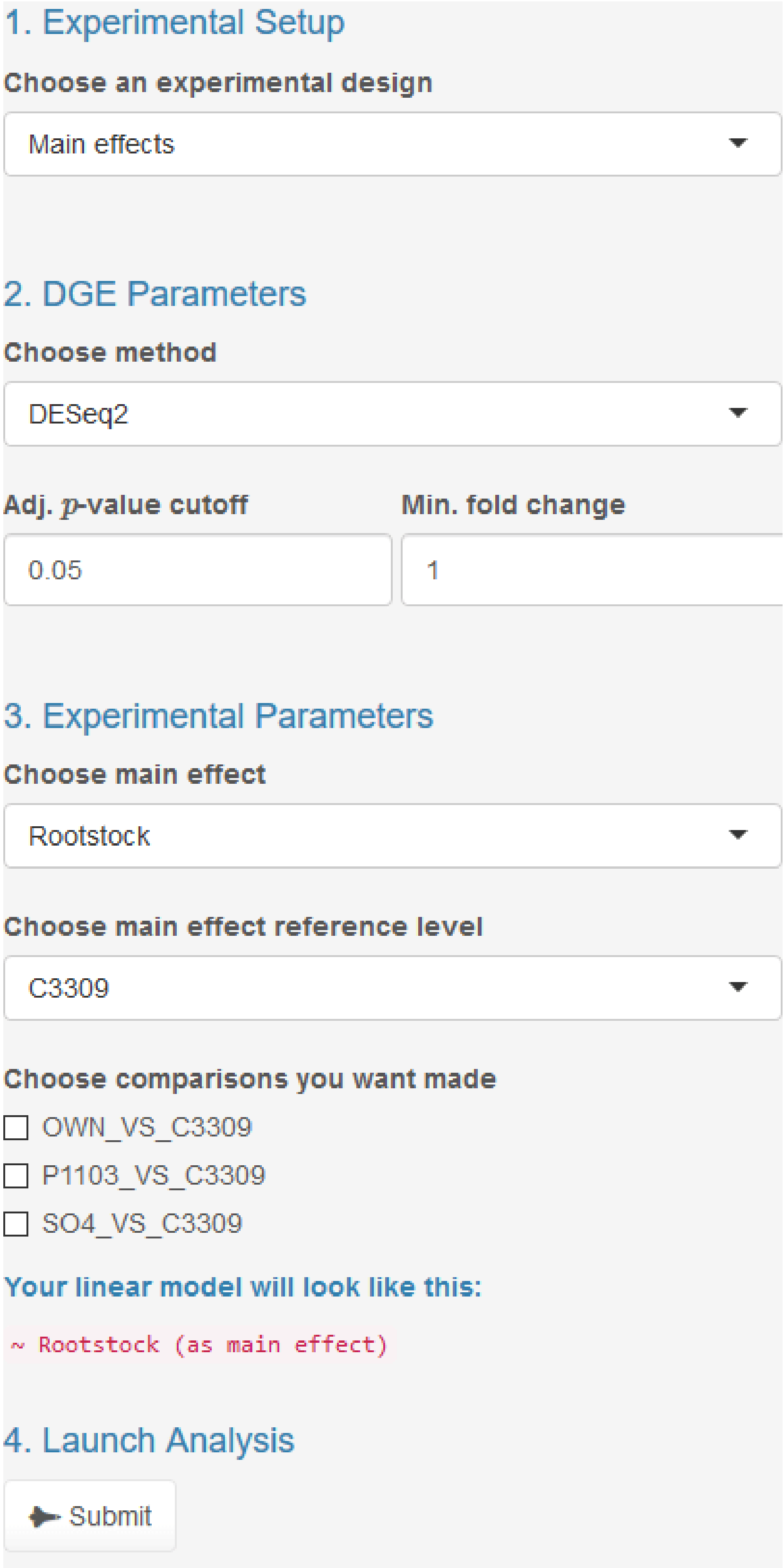
1. Experimental Setup
Choose an experimental design
Main effects
2. DGE Parameters
Choose method
DESeq2
Adj. p-value cutoff
Min. fold change
0.05
1
3. Experimental Parameters
Choose main effect
Rootstock
Choose main effect reference level
C3309
Choose comparisons you want made
OWN_VS_C3309
P1103_VS_C3309
SO4_VS_C3309
Your linear model will look like this:
~ Rootstock (as main effect)
4. Launch Analysis
Submit

6. `Main effects with grouping factors` design allows for a more detailed main effects testing to be performed. This design tests the indicated main effect when the data is subset by a user-indicated factor. Requirements for this design are indication of main effect to test, factor for grouping, and grouping factor level for which to provide results. As with the main effects design, the user also specifies a main effect reference level for lfc and visualizations. *In the example data, Rootstock is selected as the grouping factor, with C3309 selected as the grouping factor level for which the results table will be based on. The Block factor is selected to test the main effect, with B13 selected as the main effect reference level. This analysis will indicate which genes are differentially expressed when account for Block only for the C3309 samples:*

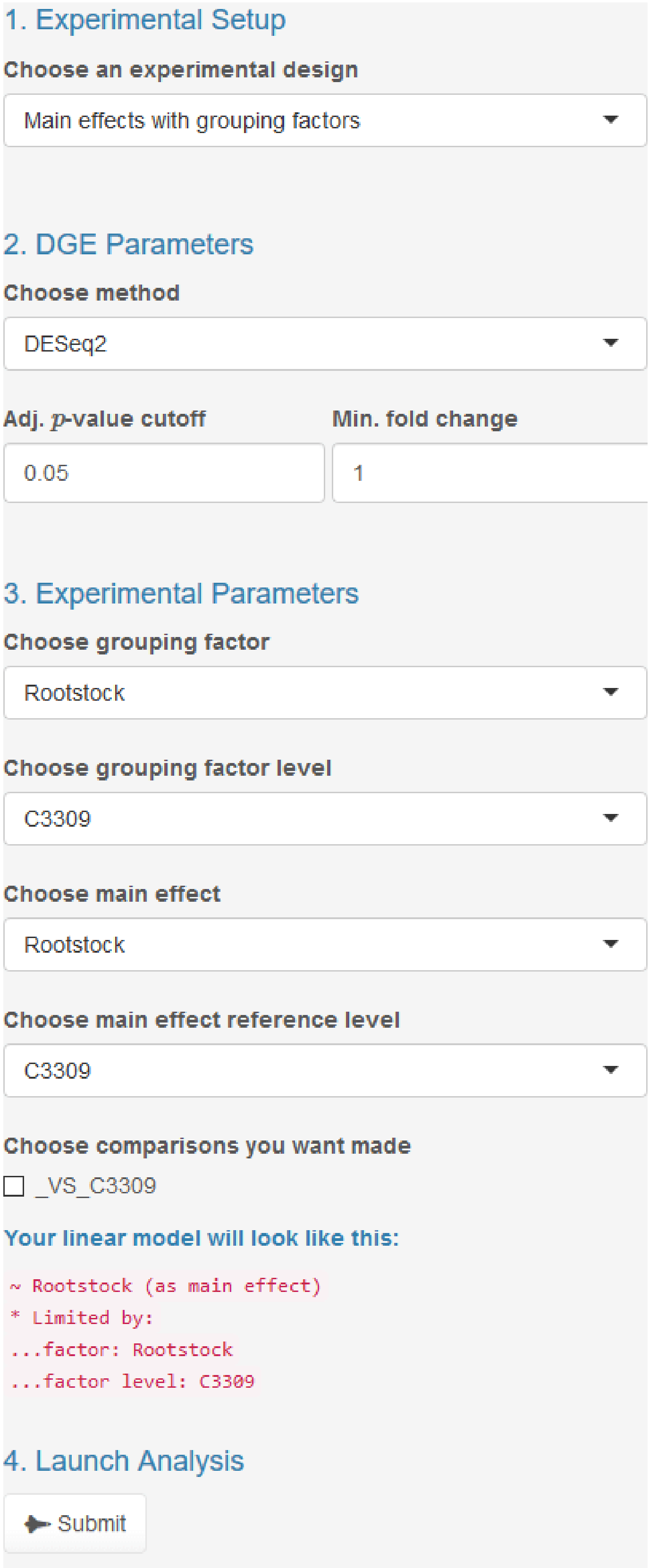
1. Experimental Setup
Choose an experimental design
Main effects with grouping factors
2. DGE Parameters
Choose method
DESeq2
Adj. p-value cutoff
Min. fold change
0.05
1
3. Experimental Parameters
Choose grouping factor
Rootstock
Choose grouping factor level
C3309
Choose main effect
Rootstock
Choose main effect reference level
C3309
Choose comparisons you want made
_VS_C3309
Your linear model will look like this:
~ Rootstock (as main effect)
* Limited by:
...factor: Rootstock
...factor level: C3309
4. Launch Analysis
Submit

7. Finally, `Custom design` provides advanced users with a method for indicating their own design matrix. This method is provided so that users can perform more intricate DGE analyses beyond what IRIS-DGE already provides as experimental design options. It is only recommended that users with advanced knowledge of the modeling process to use this option, as the required design matrix may not be intuitive to use appropriately:

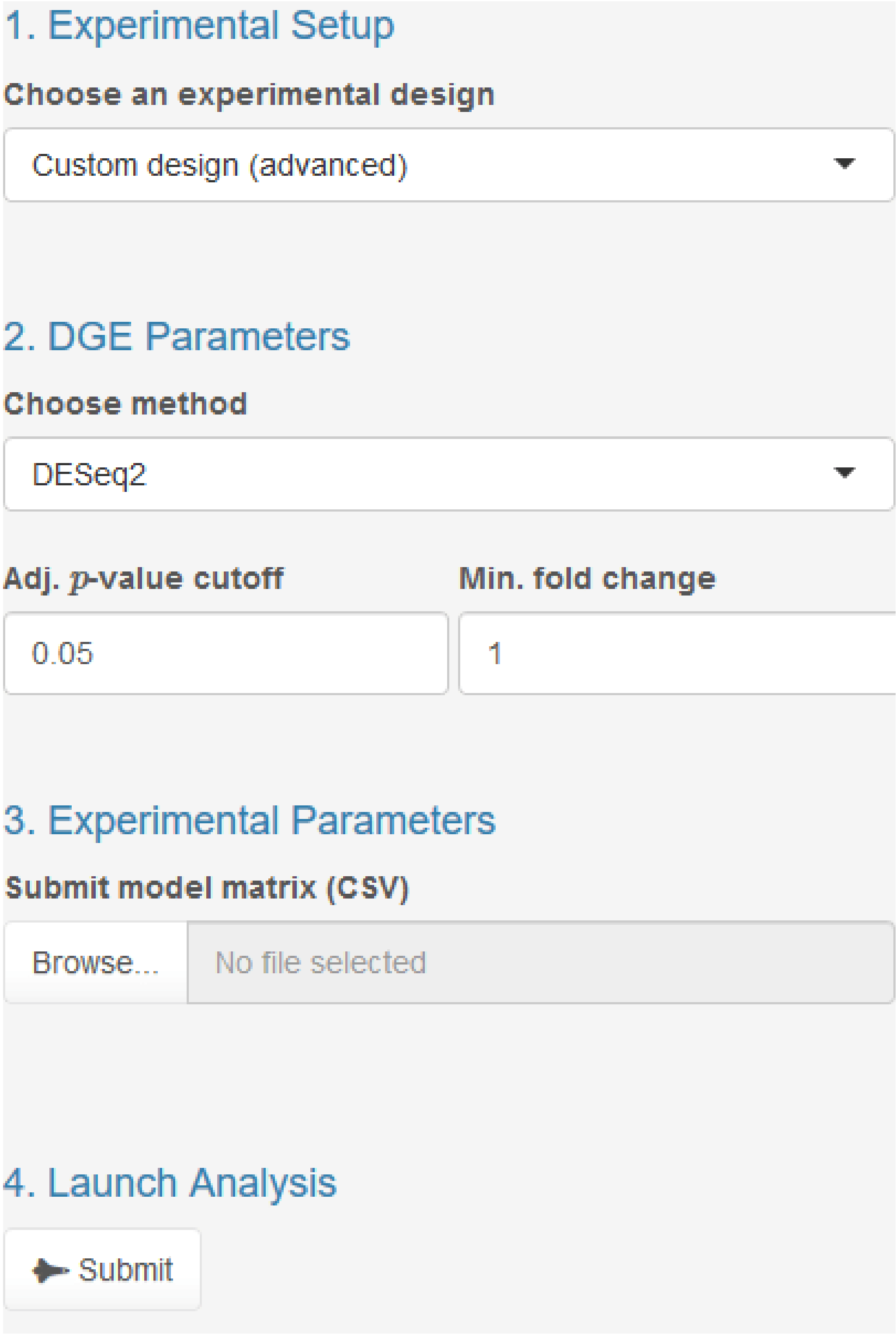
1. Experimental Setup
Choose an experimental design
Custom design (advanced)
2. DGE Parameters
Choose method
DESeq2
Adj. p-value cutoff
Min. fold change
0.05
1
3. Experimental Parameters
Submit model matrix (CSV)
Browse...
No file selected
4. Launch Analysis
Submit

#### S5.4.2: Setting up an experiment

1. Following experimental design selection, the user can then select the desired tool for performing differential gene expression analysis. This will be found under `2. DGE Parameters`. Options for this purpose are `DESeq2`, `edgeR`, and `limma-voom`. If the user selects one of the main effects design options, the only options for DGE tools are `DESeq2` and `edgeR`:

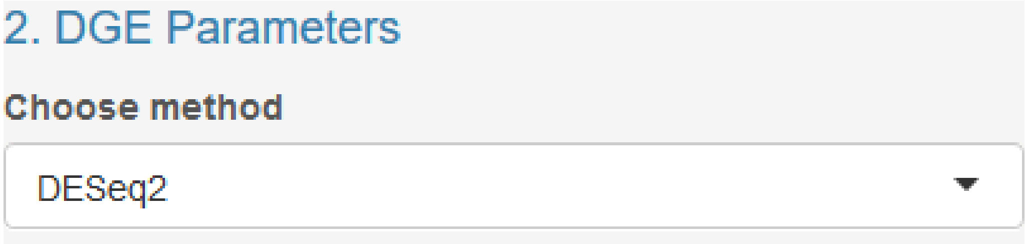

2. Also under `2. DGE Parameters`, adjusted $p$-value and log-fold change cutoffs can be specified to filter the results provided. The default values are 0.05 for adjusted $p$-value and 1 for the minimum log-fold change. To provide all gene comparisons, select 1 for adjusted $p$-value cutoff and 0 for the minimum log-fold change:

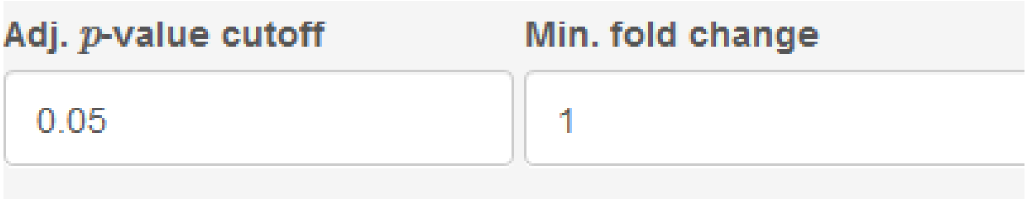

3. Upon indication of the experimental design, DGE tool and cutoffs, the parameters and comparisons for the specified analysis must be provided. Such parameters can be found under `3. Experimental Parameters`. Depending on the selected experimental design, the required parameters and comparisons will vary:

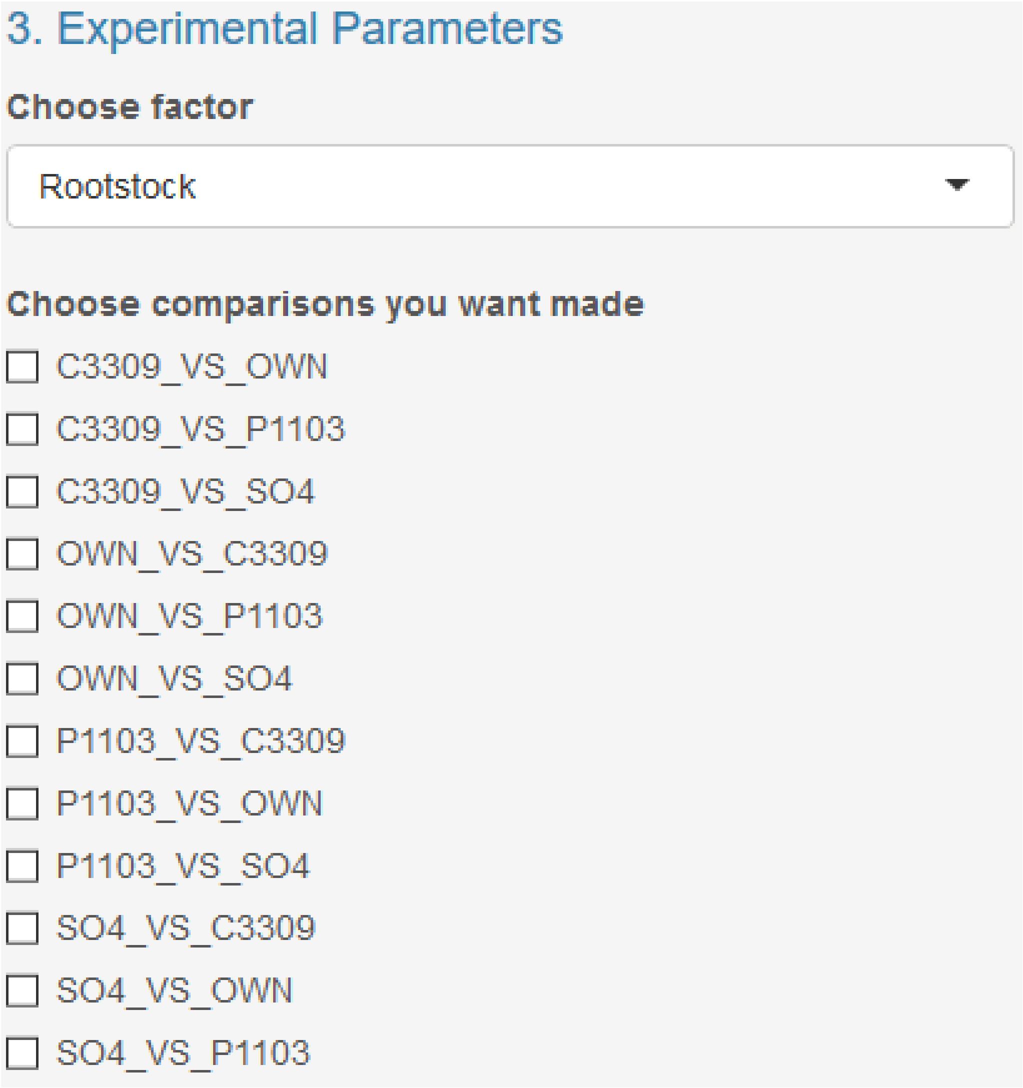

4. Launching the analysis will perform the DGE analysis and provide results. This is found under `4. Launch Analysis`. **Note:** Be mindful that this process may take more time than previous steps. Adjusting the cutoff values after submission will automatically update the generated visualizations and results table:

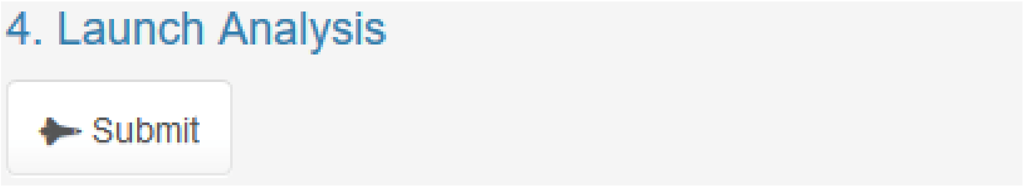

5. If you haven't crashed your computer or our server, the main page will now be populated with data in the two subtabs: `Overview` and `Plots`.

6. The `Overview` subtab (1) will display information based on the number of differentially expressed genes (DEGs), including a table of significantly up- and down-regulated gene ID counts by selected comparisons (2) and an interactive barplot representing this data

(3). *With the example data, the results show only one gene is differentially expressed in either direction from the two-group comparison based on the Rootstock factor. This information seems to follow with the previously investigated figures showing limited clustering of samples from the PCA and MDS and high correlation values between samples:*

Overview 1 Plots

DGE Overview

Show 10 entries Search:

| 2 | Comparison | Regulation | IDs |
|---|---|---|---|
| 1 | C3309_VS_OWN | up | 0 |
| 2 | C3309_VS_P1103 | up | 0 |
| 3 | C3309_VS_SO4 | up | 0 |
| 4 | OWN_VS_P1103 | up | 1 |
| 5 | OWN_VS_SO4 | up | 0 |
| 6 | P1103_VS_SO4 | up | 0 |
| 7 | C3309_VS_OWN | down | 0 |
| 8 | C3309_VS_P1103 | down | 0 |
| 9 | C3309_VS_SO4 | down | 0 |
| 10 | OWN_VS_P1103 | down | 0 |

Showing 1 to 10 of 12 entries Previous 1 2 Next

Download Table

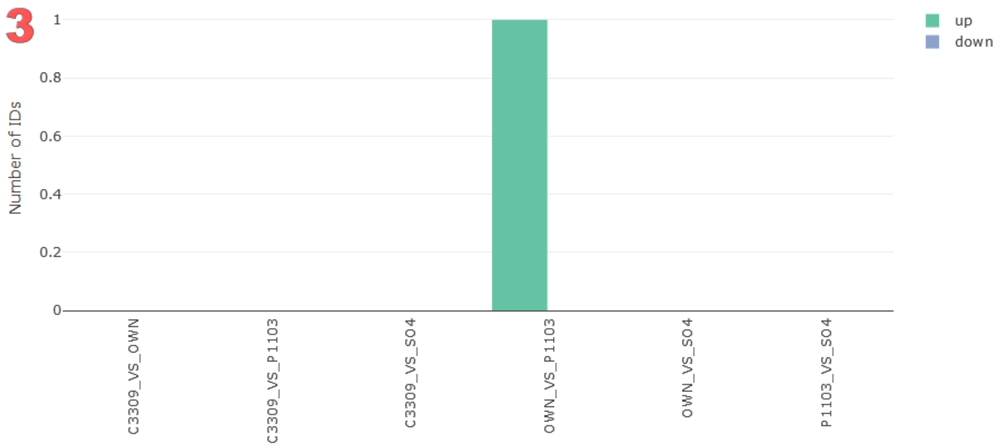

Download Static R Plot (PDF) Download Static R Plot (PNG)

7. The `Plots` subtab (1) will provides interactive visualizations based on the differential expression analysis as well as the actual results file from the previously-selected tool based on the selected comparison. The MA plot (2) shows the transformed log-fold change compared to the transformed base mean for each gene ID. Specific points can be selected, highlighting both the point and the corresponding row in the results table found below the figure. Additionally, the gene ID can be selected from the results table, which will highlight the location of that gene ID on the figure:

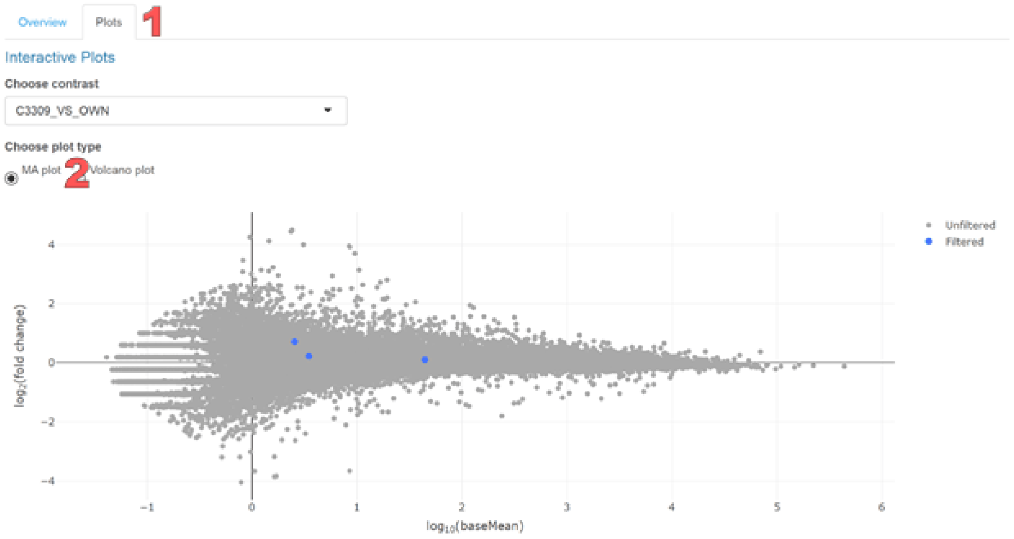

8. The Volcano plot shows a comparison of the transformed $p$-value compared with the transformed log-fold change. As with the MA plot, the Volcano plot is interactively connected with the results table found below the figure:

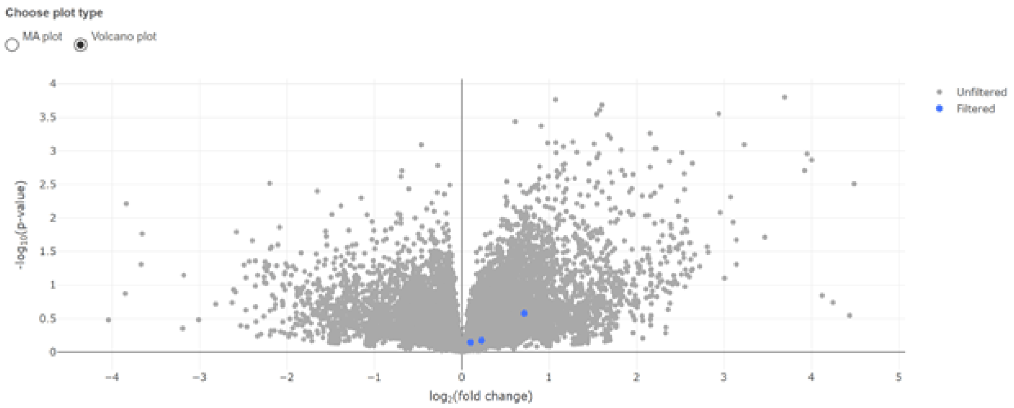

9. Finally, the results table is generated in coordination with the above plots and displays the output file of the selected tool's differential gene expression analysis. This table can be sorted increasingly or decreasingly by any of column. The integrated search feature also allows for specific gene IDs to be found in the table. This results file can be exported in a filtered or unfiltered format. *With the example data, sorting by adjusted $p$-value (padj) shows that there are no differentially expressed genes for this specific comparison (C3309 vs. OWN):*

Show 10 entries

Search:

| | id | baseMean | log2FoldChange | lfcSE | stat | pvalue | padj |
|---|---|---|---|---|---|---|---|
| 1 | gene10009 | 2.542 | 0.715 | 0.642 | 1.114 | 0.265 | 0.996 |
| 2 | gene10112 | 3.473 | 0.224 | 0.53 | 0.423 | 0.672 | 0.996 |
| 3 | gene10171 | 44.344 | 0.1 | 0.274 | 0.365 | 0.715 | 0.996 |
| 4 | gene10243 | 0.339 | -0.881 | 2.219 | -0.397 | 0.691 | 0.996 |
| 5 | gene10254 | 198.413 | 0.156 | 0.152 | 1.026 | 0.305 | 0.996 |
| 6 | gene10280 | 0.605 | 0.866 | 1.393 | 0.622 | 0.534 | 0.996 |
| 7 | gene10293 | 3.615 | -0.331 | 0.633 | -0.523 | 0.601 | 0.996 |
| 8 | gene10345 | 0.546 | -0.258 | 1.269 | -0.203 | 0.839 | 0.996 |
| 9 | gene10357 | 0.068 | 0.183 | 4.224 | 0.043 | 0.965 | 0.996 |
| 10 | gene10360 | 17.14 | -0.445 | 0.324 | -1.373 | 0.17 | 0.996 |

Showing 1 to 10 of 27,026 entries

Previous 1 2 3 4 5 … 2703 Next

Download Filtered Data Download All Data
